# HIV-1 promotes loss of CD4^+^ T cell stemness and enrichment of effector-like states via Vpr-mediated TCF7 degradation

**DOI:** 10.64898/2026.07.31.741970

**Authors:** Johanna Leyens, Anthea Darius, Carlos Alberto Vanegas-Torres, Brigitta Maurer, Ramona Businger, Nicole Wölfle, Praveen Baskaran, Stefanie M. Hauck, Michael Schindler

## Abstract

HIV-1 Vpr is abundantly packaged into virions and remodels host cells immediately after entry. Here, using high-efficiency HIV-1 infection protocols and unbiased proteomics in primary CD4+ T cells, we identify the T cell fate regulator TCF7 (TCF-1) as a previously unrecognized Vpr target. Virion-delivered Vpr rapidly depleted TCF7 in both resting and activated CD4+ T cells, which was a conserved activity of diverse Vpr proteins. The activity was independent of canonical Vpr substrate engagement but resulted terminally in proteasomal degradation of TCF7. TCF7 suppressed HIV-1 production and Env incorporation, whereas its depletion promoted loss of stem-like properties and differentiation toward effector phenotypes. Accordingly, effector T cell differentiation states are associated with productive HIV-1 infection and reduced TCF7 abundance. Thus, Vpr-mediated TCF7 depletion couples enhanced viral fitness to reprogramming of CD4+ T cell identity, generating permissive differentiated cells while impairing the maintenance of effective antiviral immunity via reduced T cell stemness.

## INTRODUCTION

Vpr is a small accessory protein, often dispensable for replication in immortalized cell lines but important in a physiological context. It is found in lentiviruses, approximately 96 amino acids in length, with a molecular weight of 12–14 kDa. Due to its conserved presence within the HIV-1 genome and its critical role in the pathogenicity of both HIV-1 and SIV *in vivo*, Vpr is believed to manipulate host cells to allow immune evasion and facilitate viral replication [1]. However, on a molecular level, the precise mechanisms by which Vpr contributes to viral pathogenesis and elevated viral loads remain elusive [2], even though growing evidence suggests a role for Vpr in priming resting T cells for productive infection [3–5].

Vpr can be considered an HIV-1 encoded proteolysis targeting chimera (PROTAC). By this, Vpr specifically remodels the host cell proteome. How Vpr achieves profound deregulation of cellular proteins by targeted degradation of host cell factors is well understood. Functioning as an adaptor protein, Vpr facilitates the recruitment of specific target proteins to DCAF1 (DDB1- and CUL4- Associated Factor 1), also known as VPRBP (Vpr-binding protein) [6–8]. DCAF1 serves as a substrate recognition receptor, linking Vpr-associated targets to the CUL4A E3 ubiquitin ligase complex, ultimately leading to their proteasomal degradation. To achieve this, Vpr binds to DCAF1, an interaction that can specifically be disrupted via mutation of a glutamine at position 65 (Q65) in Vpr [6, 9]. Furthermore, Vpr recruits certain cellular factors, likely via multiple domains, and channels them for degradation. Of note, depletion of host factors targeted for degradation by Vpr through the DCAF1 pathway has been shown to result in G2 arrest. This effect has been observed in various cell lines, including HEK293T, HeLa cells, and immortalized CD4^+^ T cell lines such as Jurkat and SupT1 [2, 7, 10]. Examples of factors proposed to be involved in this process include MCM10 [11], the exonuclease EXO1 [12], and CCDC137 [13]. These findings led to the widely accepted hypothesis that Vpr selectively recruits and degrades host proteins via the DCAF1-E3-CUL4A axis to mediate G2 arrest in infected cells [14], even though the biological relevance of this function is unclear and Vpr does not seem to induce a G2 arrest in primary CD4^+^ T cells [2, 4]. Altogether, to better understand and potentially find novel as yet-undefined functionalities of the Vpr-DCAF1-Cul4A axis, efforts have been undertaken to comprehensively map Vpr-induced proteomic changes on a cellular level. To achieve this, Vpr-interacting proteins were purified and characterized from monocytic and T cell lines identifying DNA damage response associated proteins such as MUS81 and EME1 [15], or the HLTF helicase [16]. Furthermore, cell-based proteomics found EXO1 as a factor depleted in Vpr-overexpressing cells [12]. The provisional final assessment came from the landmark paper from the Lehner lab, which performed comprehensive multi-level proteomics on a T cell line to identify Vpr-interacting as well as Vpr-depleted host cell factors [17]. The analyses led to a final line-up of at least 38 Vpr-depleted factors, the majority of them involved in cell cycle and DNA-damage repair, among them the previously established DCAF1, MUS81, ZGPAT and HLTF [17].

Immortalized cell lines often have defects in antiviral and antitumor interferon-signaling pathways as well as cell cycle control and DNA-damage response [18]. We hence hypothesized that Vpr-induced proteomic changes in primary CD4^+^ T cells might differ from those observed in cell lines and further reveal HIV-1 factors targeted for proteasomal degradation. Therefore, we used an established high efficiency HIV-1 infection protocol in primary CD4^+^ T cells [4], which were then subjected to non-targeted proteomics. This approach revealed depletion of known host cell factors from HIV-1 infected CD4^+^ T cells in a Vpr-dependent manner. We further found previously unprecedented protein depletions attributable to Vpr, with the transcription factor TCF7 representing one prominent hit.

TCF7, together with the Wnt-effector β-Catenin canonically regulates genes involved in cell survival, proliferation and differentiation [19–22]. Multiple isoforms of this T cell–specific transcription factor have been identified and can be categorized into β-catenin binding isoforms, which become active upon β-catenin binding, as well as truncated isoforms which do not comprise a β-catenin binding domain and function as transcription repressors. In mature CD8^+^ T cells, TCF7 is particularly important for preserving less differentiated cellular states, supporting stemness-associated programs and sustaining self-renewal, highlighting TCF7 to be considered as an important factor to improve CAR T cell-based therapies [23]. Similarly, in CD4^+^ T cells, TCF7 is supposed to be critical for preserving “stemness”, allowing T cells to differentiate into effector cells [24–27]. Furthermore, TCF7 influences polarization of conventional CD4^+^ T cell subsets. High TCF7 levels favor Tfh over Th1 differentiation, repressing IFNγ expression and inducing Th2 differentiation by promoting GATA3 expression [28–32]. In contrast, reduced TCF7 expression might destabilize the Th17/Treg balance [20]. Herein, we identified that TCF7 is degraded in CD4^+^ T cells upon HIV-1 infection in a Vpr-dependent manner. We characterized the mechanism of Vpr-driven TCF7 depletion and the functionality of various Vpr alleles and mutants in this process. We further investigate the functional relevance of this phenomenon for HIV-1 replication. Altogether, we identify TCF7 as a key regulator of T cell differentiation that HIV-1 manipulates in primary CD4^+^ T cells.

## RESULTS

### Identification of HIV-1 and Vpr-deregulated proteins in primary CD4^+^ T cells

To enable unbiased, non-targeted proteomics, we used our high-titer infection protocol for primary CD4^+^ T cells [4]. This reduces background signal from non-infected cells and improves the signal-to-noise ratio. PHA/IL-2 pre-stimulated primary CD4^+^ T cells were incubated with sucrose-purified viral supernatants and spinoculated for infection. HIV-1- or ΔVpr-infected samples were then cultured for 24 h and 48 h. Aliquots were stained for cell-associated HIV-1 p24 to assess infection rates by flow cytometry (Fig. 1A), while the remaining cells were lysed and subjected to untargeted whole-cell proteomics. To assess Vpr-attributable changes, time dependence, and potential entry-dependent differences, we included VSV-G-pseudotyped HIV-1. In the four donors used for proteomics, infection rates ranged from 70-90% and were slightly higher at 48 hpi and with VSV-G entry as compared to HIV-1 GP120 (Fig. 1A). We calculated proteome changes based on relative protein abundances across the conditions. Overall, we detected at least one peptide annotated to 7587 human proteins (Supplemental Table S1). Mean values of the relative arbitrary protein abundances from the four donors were used to calculate fold changes in pairwise comparisons across all conditions, which were assessed for statistical significance using a standard T-test. We then generated volcano plots depicting significant fold changes in protein abundances for all pairwise comparisons (Supplemental Table S1). To make the data transparent and accessible, we prepared an interactive, automated table that facilitates rapid retrieval of all analyzed proteins (Supplemental Table 2). To identify the most pronounced and HIV-1-relevant changes in the cellular proteome, we first analyzed HIV-1-induced changes relative to non-infected cells (mock) at 48 hpi (Fig. 1B). As expected, and as an important internal control, HIV-1 gene products Gag, Gagpol, Env, and Nef were strongly enriched. Focusing on proteins that were less abundant in HIV-1-infected cells, we identified CD4 as a target for directed degradation by the viral proteins Nef, Vpu, and Env [33, 34], as well as known Vif targets belonging to the APOBEC family and PP2A phosphatases [35, 36]. Among several other proteins that were reduced at least 2-fold, we identified only three that were also hits when comparing HIV-1-infected CD4^+^ T cells to the ΔVpr-infected samples at 48 h (Fig. 1C and D). These were ACTG1 (Gamma-actin), DGKB (Diacylglycerol kinase beta) and TCF7 (Transcription factor 7). Of these, DGKB and TCF7 showed consistent reduction independent of viral entry at 48 hpi (Fig. 1E). Given that TCF7 emerges as a potential key transcription factor involved in T cell development and orchestration of immune responses [20, 28, 30, 37, 38], we focused on this factor as a potential new Vpr target in HIV-1-infected primary CD4^+^ T cells. Accordingly, we set out to confirm Vpr-dependent reduction of TCF7 upon HIV-1 infection by repeating the high-titer infection protocol and assessing TCF7 levels by Western blot analysis (Fig. 1F). Indeed, TCF7 seems nearly completely depleted in CD4^+^ T cells infected up to 80% with HIV-1. Conversely, TCF7 levels were similar to those of non-infected cells upon infection with HIV-1 carrying an inactivating deletion in the *vpr* open reading frame (ORF). Altogether, using non-targeted proteomics, we identify TCF7 as a host factor depleted in HIV-1-infected primary CD4^+^ T cells in a Vpr-dependent manner.

**Figure 1.**
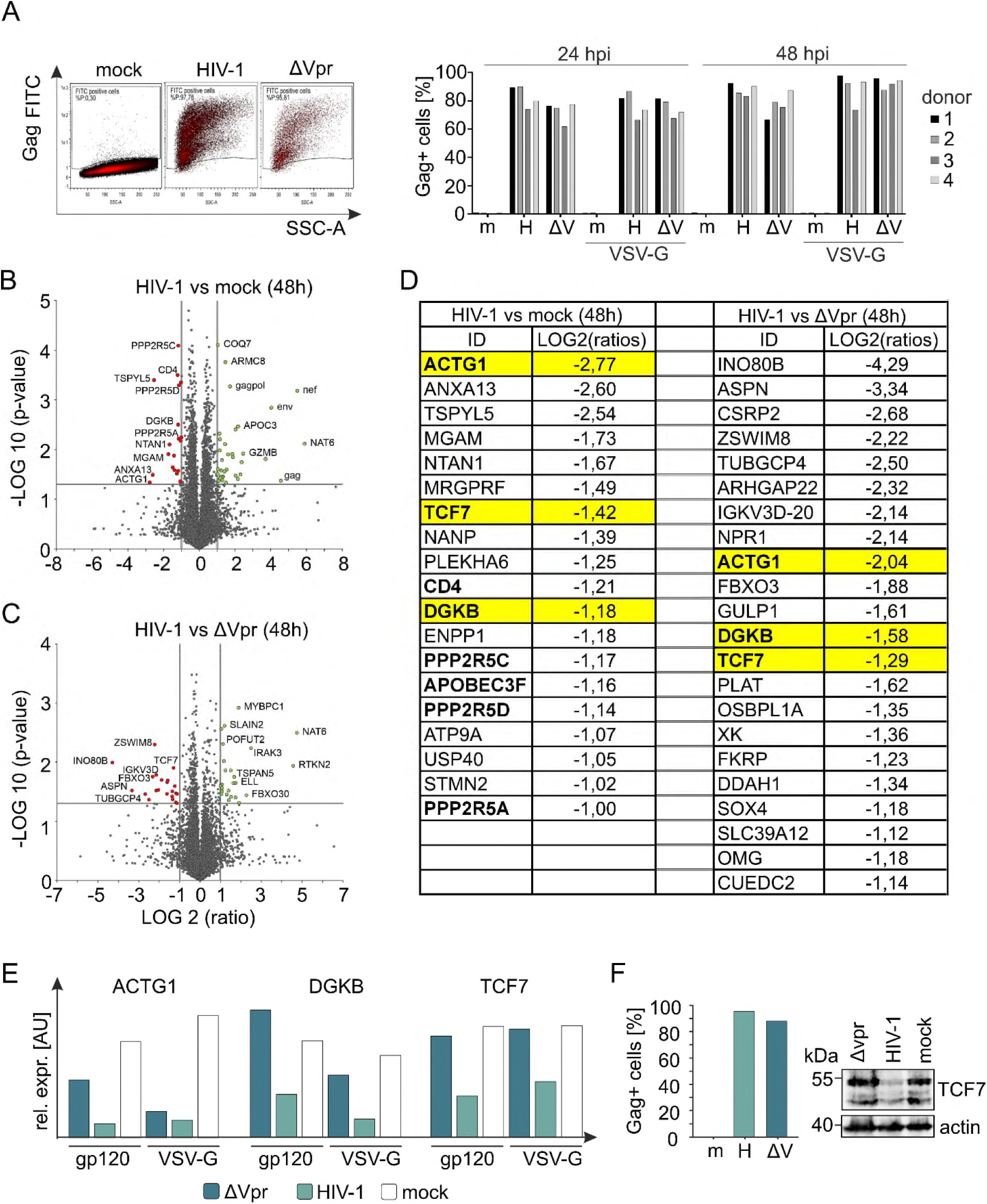
Non-targeted proteomic analysis of HIV-1-infected primary CD4^+^ T cells. (A) Primary CD4^+^ T cells were isolated from four healthy donors and infected with HIV-1 (H) or HIV-1 ΔVpr (ΔV), either VSV-G-pseudotyped or non-pseudotyped. At 24 and 48 h post-infection (hpi), cells were harvested for intracellular p24 staining and lysed for non-targeted proteomic analysis. (B and C) Volcano plots showing the log2-fold modulation and corresponding p values of protein abundance in (B) HIV-1-infected compared with mock-infected primary CD4+ T cells or (C) HIV-1-infected compared with HIV-1 ΔVpr-infected primary CD4^+^ T cells at 48 hpi. (D) Table presenting host-cell proteins with a significant reduction in abundance of more than 1 log2-fold when comparing HIV-1-infected with mock-infected or HIV-1 ΔVpr-infected primary CD4^+^ T cells at 48 hpi (p < 0.05). Proteins that were significantly modulated in both comparisons are highlighted in yellow. (E) Primary data from the non-targeted proteomic analysis showing the relative expression of the three candidate factors ACTG1, DGKB, and TCF7 in arbitrary units (AU) at 48 hpi. Mean arbitrary units from the four donors analyzed are shown. Data were obtained from the interactive Supplementary Table 2. (F) Validation of the HIV-1 Vpr-mediated reduction in TCF7 abundance in HIV-1-infected primary CD4^+^ T cells at 48 hpi. Cells were infected using the high- efficiency infection protocol and analyzed by intracellular HIV-1 Gag staining using flow cytometry and by Western blot analysis of TCF7 and actin protein levels, with the latter being the loading control.

### Virion-delivered Vpr depletes TCF7 early after infection of primary CD4^+^ T cells and TCF7 depletion is a conserved function of primary patient-derived HIV-1 alleles

To further characterize the kinetics and modes of TCF7 reduction by Vpr, we returned to the proteomic dataset and found that TCF7 was already reduced by 24 hpi in an entry-independent fashion in PHA/IL-2–stimulated primary CD4^+^ T cells, with a more pronounced loss by 48 hpi (Fig. 2A and Supplemental Table S2). PHA-based activation is a polyclonal mitogen that activates T cells independent of antigen-specific TCR engagement. Because TCF7 expression itself is known to be tightly regulated by TCR signal strength, we asked whether TCR-activation via CD3/CD28 or IL-7/IL-15 cytokine driven activation of CD4^+^ T cells display the same phenotype as in T cells stimulated with PHA/IL2 and observed a significant reduction in both the proportion of TCF7 positive cells as well as in TCF7 levels (measured by flow cytometry) at 48 hours post-infection (Fig. 2B-2D). Notably, this reduction was not only evident in cells scoring HIV-1 Gag-positive, but also in Gag-negative, i.e. uninfected bystander cells (Fig. 2D). Given that TCF7 protein levels were already reduced at 24 hours post-infection, we next examined the dynamics of TCF7 reduction, sampling HIV-1-infected primary CD4^+^ T cells between 4 and 24 hpi (Fig. 2E). This revealed Vpr-dependent TCF7 depletion already at 4 hpi, indicating that TCF7 loss is induced post-entry by virion-delivered Vpr directly. Of note, HIV-1 expressing a mutated form of Vpr (R80A) that is compromised in its ability to trigger degradation of host cell factors via the DCAF1 E3 ubiquitin ligase [39] was also attenuated in its ability to reduce TCF7 (Fig. 2E). Since virion-delivered Vpr might be sufficient to reduce TCF7, we hypothesized that HIV-1 not only degrades TCF7 in prestimulated primary CD4^+^ T cells, where efficient integration and viral gene expression are facilitated, but might also degrade TCF7 in unstimulated primary CD4^+^ T cells. Indeed, similar to infection of PHA/IL2 pre-treated CD4^+^ T cells, we observed robust Vpr-dependent TCF7 depletion in HIV-1-infected, unstimulated and hence resting primary CD4^+^ T cells (Fig. 2F).

**Figure 2.**
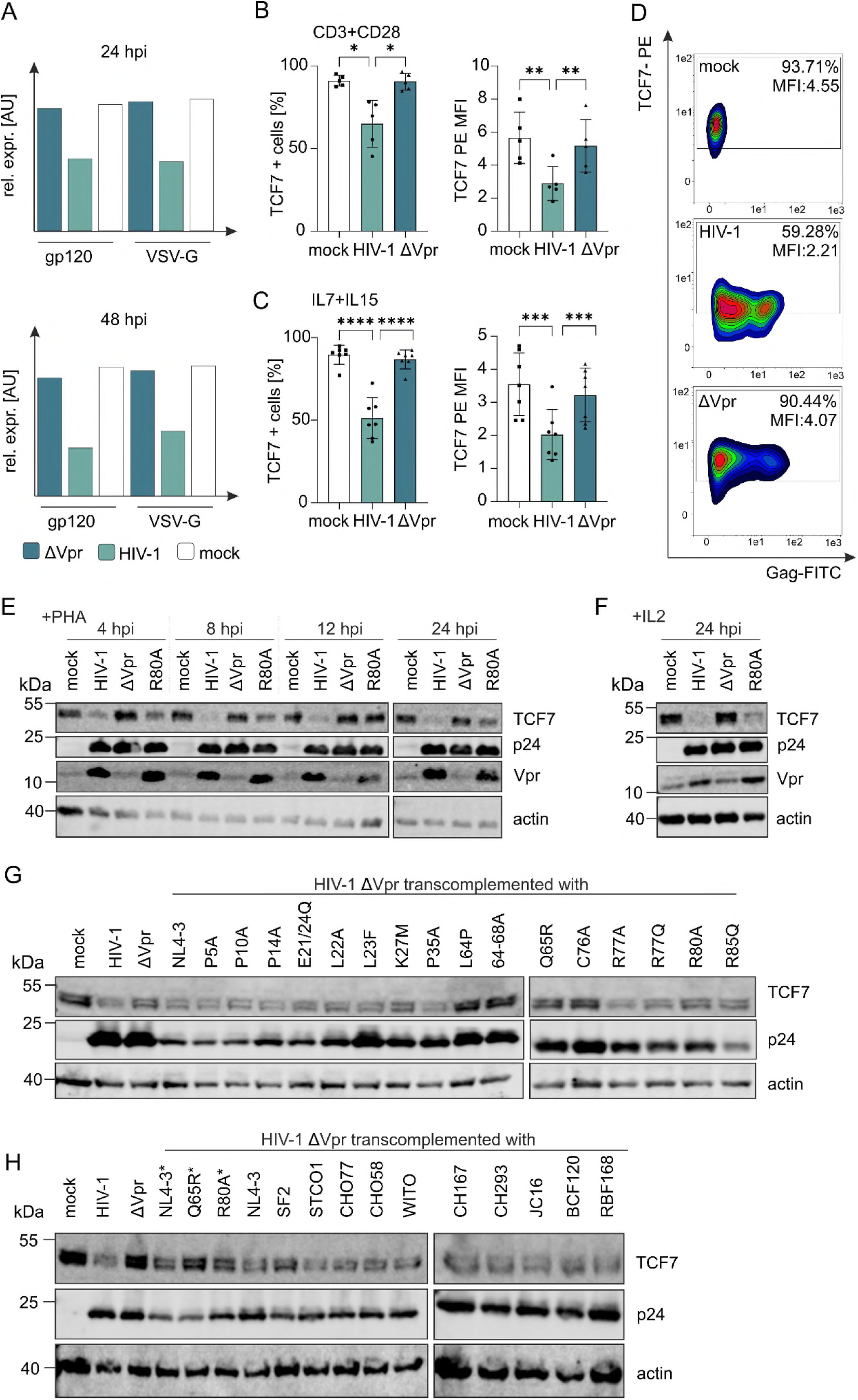
Dynamics and determinants of the HIV-1 Vpr-mediated reduction in TCF7. (A) Primary data from the non-targeted proteomic analysis showing relative TCF7 abundance in arbitrary units (AU) at 24 and 48 hpi. Mean arbitrary units from the four donors analyzed are shown. Data were obtained from the interactive Supplementary Table 2. (B and C) Primary CD4^+^ T cells were TCR-stimulated using plate-bound anti-CD3 antibody (2 µg/mL) and soluble anti- CD28 antibody (1 µg/mL) in the presence of IL-2 (10 ng/mL) and infected with HIV-1 or ΔVpr. TCF7 protein expression was determined at 48 hpi by intracellular staining. (B) together with the percentage of TCF7+ cells and TCF7 median fluorescence intensity (MFI). n=5 (Mean ± SD). Statistical significance was determined using a paired one-way ANOVA with Šídák multiple- comparison correction. *p < 0.05; **p < 0.01. (C) Primary CD4^+^ T cells were stimulated with IL-7 and IL-15 (20 ng/mL each) and infected with HIV-1 or ΔVpr. TCF7 protein expression was determined at 48 hpi. The percentage of TCF7+ cells and TCF7 MFI are shown. n=7 (Mean ± SD). Statistical significance was determined using a paired one-way ANOVA with Šídák multiple- comparison correction. ***p < 0.001; ****p < 0.0001. (D) Representative density plots of TCF7 expression in relation to Gag expression in primary CD4^+^ T cells stimulated with IL-7 and IL-15 (20 ng/ml each) upon HIV-1 infection are shown. (E) Primary CD4^+^ T cells were stimulated with IL-2 (10 ng/mL) and PHA (1 µg/mL) and infected with VSV-G-pseudotyped HIV-1, ΔVpr, or HIV-1 encoding the Vpr R80A mutant. Cells were harvested at 4, 8, 12, and 24 hpi and subjected to Western blot analysis of TCF7, HIV-1 p24, and Vpr. Actin was used as a loading control. Representative results from two biological replicates are shown. (F) Resting primary CD4^+^ T cells maintained in IL-2 (10 ng/mL) were infected with HIV-1 or ΔVpr directly after isolation without prior stimulation. Cells were lysed at 24 hpi and subjected to Western blot analysis of TCF7, HIV-1 p24, and Vpr. Actin was used as a loading control. Representative results from three biological replicates are shown. (G) SupT1 CD4^+^ T cells were infected with VSV-G-pseudotyped ΔVpr particles transcomplemented in the virus-producing cells with the indicated Vpr mutant proteins. Full-length HIV-1 was included as a positive control. Cells were harvested at 24 hpi and subjected to Western blot analysis of TCF7 and HIV-1 p24. Actin was used as a loading control. Representative blot from four biological replicates is shown. (H) SupT1 CD4^+^ T cells were infected with VSV-G-pseudotyped ΔVpr particles transcomplemented in the virus-producing cells with AU1-tagged Vpr proteins derived from different primary HIV-1 isolates. Full-length HIV-1 and the non-AU1-tagged versions of NL4-3 Vpr and the Vpr mutants Q65R and R80A, indicated by asterisks, were included as controls. Cells were lysed at 24 hpi and subjected to Western blot analysis of TCF7 and HIV-1 p24. Actin was used as a loading control. Results are representative of three biological replicates.

To determine whether incoming virion-associated Vpr was sufficient to drive TCF7 reduction and to map the structural residues of Vpr required for its activity, we employed Vpr-deficient HIV-1 and transfected HEK293T producer cells to co-express Vpr or a panel of Vpr mutants in trans, allowing Vpr to be transcomplemented and packaged into viral particles. Subsequently, we infected SupT1 CD4^+^ T cells and assessed TCF7 levels by Western blot (Fig. 2G). In comparison to HIV-1 ΔVpr transcomplemented with a GFP-expression vector only, particles that incorporated HIV-1 NL4-3 Vpr clearly reduced TCF7. As expected, Vpr-mutants that are not incorporated into progeny virions (L64P and 64-68A) failed to deplete TCF7. Similarly, Vpr mutants that fail to induce G2-arrest in immortalized cells and are reported to be impaired or defective in their interaction with the DCAF1 adaptor protein complex were fully (Q65R, C76A) or partially (K27M, R80A) defective for TCF7 reduction (Fig. 2G). Furthermore, when de novo viral gene expression was inhibited by blocking either reverse transcription by treatment with Efavirenz or integration with Raltegravir, Vpr was still able to deplete TCF7 early after infection (Supplementary Fig. S1A). In conclusion, virion-delivered Vpr is sufficient to trigger TCF7 reduction involving domains necessary for Vpr interaction with the DCAF1 adaptor protein complex.

Viral factors other than Vpr could be involved in TCF7 depletion, and primary Vpr alleles might functionally differ from those of the HIV-1 NL4-3 strain thus far employed, which is a lab-adapted T cell passaged strain. We therefore tested if Vpr alone and Vpr alleles from different HIV-1 groups and subtypes induce TCF7 reduction. For this, we employed lentiviral constructs expressing Vpr only and used them to transduce the SupT1 CD4^+^ T cell line (Supplementary Fig. S1B). In addition, within the same lentiviral backbone, we used a collection of primary Vpr alleles derived from HIV-1 subtypes B, C, D, and H, as well as HIV-1 groups N, O, and P. Furthermore, an extended version of subtype D Vpr (D_lo) that fails to induce G2 arrest in cell lines, as well as a mutant of the HIV-1 P Vpr (Q65L) that does not interact with DCAF1, were included. In SupT1 CD4^+^ T cells, even though infection efficiency varied based on the levels of cells-associated p24 that is delivered alongside the lentiviral particle, all primary HIV-1 M subtype Vpr alleles as well as the ones derived from HIV-1 groups N, O, and P efficiently depleted TCF7 (Fig S1B). Consistent with our data using the Vpr NL4-3 mutants (Fig. 2G), alleles defective in inducing cellular G2 arrest, i.e., HIV-1 subtype D_lo and HIV-1 P_Q65L, also failed to reduce TCF7. We extended this analysis to a variety of patient-derived HIV-1 transmitted/founder and chronic strains and transcomplemented the respective Vpr alleles by co-transfecting expression plasmids into the producer cells. Upon infection of SupT1 CD4^+^ T cells, all Vpr alleles caused TCF7 depletion (Fig. 2H). In conclusion, interference with steady-state TCF7 protein levels is highly conserved across HIV-1 subtypes and represents a common feature of primary, patient-derived Vpr alleles.

### Proteasomal degradation of TCF7 is not mediated by canonical Vpr engagement

Vpr mutants that are impaired or defective in their ability to hijack DCAF1, including Vpr Q65R/L, C76A, K27M or R80A, failed to reduce TCF7 in SupT1 cells (Fig. 2G and Fig. S1B). We therefore hypothesized that TCF7 might be degraded via the proteasomal pathway and that Vpr acts as an adaptor to direct TCF7 for proteasomal degradation via the Vpr-associated DCAF1 E3 ubiquitin ligase complex [39–42]. Accordingly, we treated HIV-1-infected primary CD4^+^ T cells with the proteasome inhibitor MG132 or an inhibitor of lysosomal degradation, bafilomycin A1 (BafA1), as a control (Fig. 3A). In line with a model of proteasomal TCF7 degradation via Vpr, MG132 treatment restored TCF7 levels in comparison to non-infected cells or CD4^+^ T cells infected with the HIV-1 ΔVpr variant. In contrast, BafA1 did not have any effect on Vpr-induced TCF7 degradation (Fig. 3A). We furthermore analyzed TCF7 mRNA levels via qRT-PCR in HIV-1- infected primary CD4^+^ T cells and did not detect any changes in TCF7 transcripts, neither in HIV- 1 versus non-infected cells nor as a result of functional Vpr-expression (Fig. 3B). Furthermore, we aimed to investigate whether Vpr directly interacts with TCF7. For this, HEK293T cells were transfected to express HA-tagged TCF7, harboring a β-catenin binding site, and HIV-1 NL4-3 Vpr and the Vpr Q65R mutant. To stabilize the potential complex and prevent TCF7 degradation, we also employed the proteasome inhibitor MG132. TCF7 and its interaction partners were then purified by immunoprecipitation against the HA tag, and both input and pull-down fractions were analyzed by Western blot (Fig. 3C). Notably, although the TCF7-HA pull-down efficiently co-precipitated β-catenin, an endogenous TCF7-binding partner expressed in HEK293T cells [43–45], there was no direct interaction between TCF7 and Vpr detectable, regardless of MG132 treatment. Furthermore, TCF7-HA did not co-immunoprecipitate with DCAF1, independent of Vpr expression. Together, these results indicate that although Vpr targets TCF7 for proteasomal degradation, it does not directly interact with TCF7, nor is TCF7 a canonical DCAF1-binding partner, suggesting an indirect degradation mechanism that remains to be defined.

**Figure 3.**
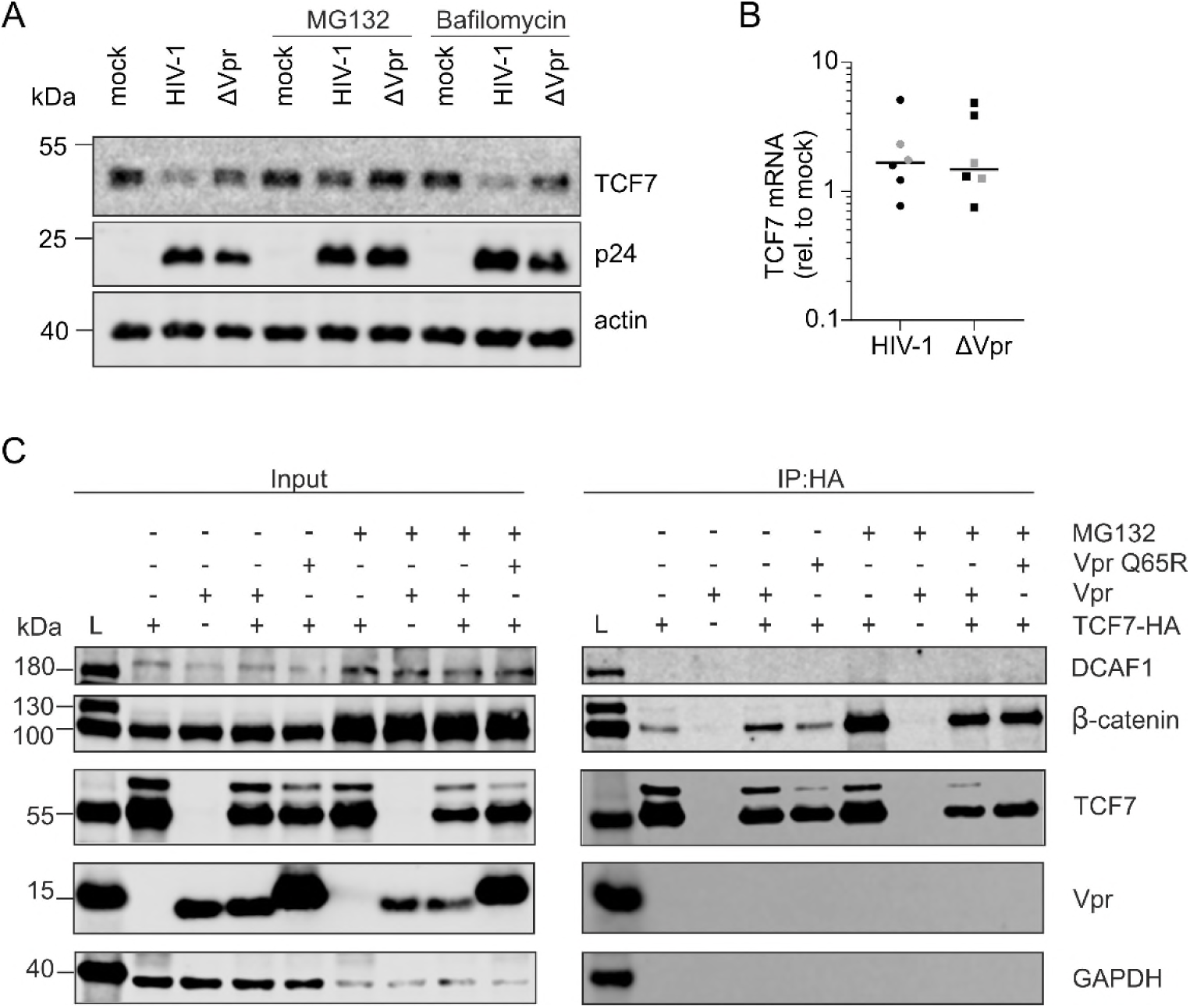
Mechanism of Vpr-mediated TCF7 degradation. (A) Primary CD4^+^ T cells prestimulated with PHA (1 µg/mL) and IL-2 (10 ng/mL) were infected with VSV-G-pseudotyped HIV-1 or ΔVpr. At 2 hpi, the proteasome inhibitor MG132 (1 µM), the lysosomal inhibitor bafilomycin A1 (500 nM), or DMSO as a vehicle control was added. Cells were lysed at 12 hpi and subjected to Western blot analysis of TCF7 and HIV-1 p24. Actin was used as a loading control. Representative results from three biological replicates are shown. (B) Primary CD4^+^ T cells prestimulated with PHA (1 µg/mL) and IL-2 (10 ng/mL) were infected with VSV-G- pseudotyped (gray symbols) or non-pseudotyped (black symbols) HIV-1 or ΔVpr. At 48 hpi, cells were harvested for RNA isolation, and TCF7 mRNA abundance was determined by RT-qPCR using GAPDH as the housekeeping gene. Relative TCF7 mRNA abundance, calculated using the 2−ΔΔCt method, is shown. n=6 (Mean). (C) Co-immunoprecipitation of HA-tagged TCF7 was performed following transfection of HEK293T cells with TCF7-HA and NL4-3 Vpr. The Vpr Q65R mutant was included as a negative control. At 6 h post-transfection, the medium was replaced with medium containing MG132 (1 µM) or DMSO as a vehicle control. Following cell lysis, TCF7 and its interacting proteins were immunoprecipitated and subjected to Western blot analysis for DCAF1, β-catenin, TCF7, Vpr, and GAPDH. β-Catenin was assessed as a positive control for efficient TCF7 immunoprecipitation. Representative blot from three biological replicates is shown.

### TCF7 interferes with infectious HIV-1 particle production

We next determined the potential impact of TCF7 on basic parameters of viral replication, i.e., particle production and release of infectious virions. Therefore, 293 T cells were transfected to produce HIV-1 NL4-3 in the presence of increasing amounts of TCF7 or mTCF7 with a mutated β-catenin binding site. Then, virus production in cell culture supernatants was analyzed by p24 ELISA (Fig. 4A). Notably, TCF7 expression led to a dose-dependent reduction in released viral particles, as evidenced by lower p24 levels. Apparently, Vpr seems not to be able to counteract this effect in 293T cells, as levels of supernatant-associated p24 were reduced, independent of Vpr expression (Fig. 4A). Furthermore, an intact β-Catenin binding site in TCF7 is essential, since the mutant version mTCF7 does not exert any negative effect on virus production and release. We also analyzed the infectivity of released viral particles. Normalized amounts of HIV-1 NL4-3 p24 or the ΔVpr variant produced in the presence of increasing TCF7 levels were used to infect LC5- RIC reporter cells [46]. Then, infection was quantified via reporter gene expression and depicted relative to infection without TCF7 co-expression in the producer cells (Fig. 4B). This revealed that TCF7 reduced the particle infectivity of HIV-1 ΔVpr, again dependent on an intact β-Catenin binding site. No differences in HIV-1 particle infectivity were detected when mTCF7 was co-expressed in the producer cells or upon functional HIV-1 Vpr expression (Fig. 4B). To gain functional insights into the potential underlying reason for reduced HIV-1 particle infectivity, we performed Western blot analysis of HEK293T HIV-1 producer cells as well as viral supernatants. This revealed that, in relation to p24, TCF7, but not mTCF7, reduced cell-associated levels of the HIV-1 envelope glycoprotein in a Vpr-dependent manner (Fig. 3C). Altogether, TCF7 containing an intact β-catenin binding site suppresses the release of infectious HIV-1 particles, indicating a TCF7-dependent antiviral effect that is partially antagonized by Vpr.

**Figure 4.**
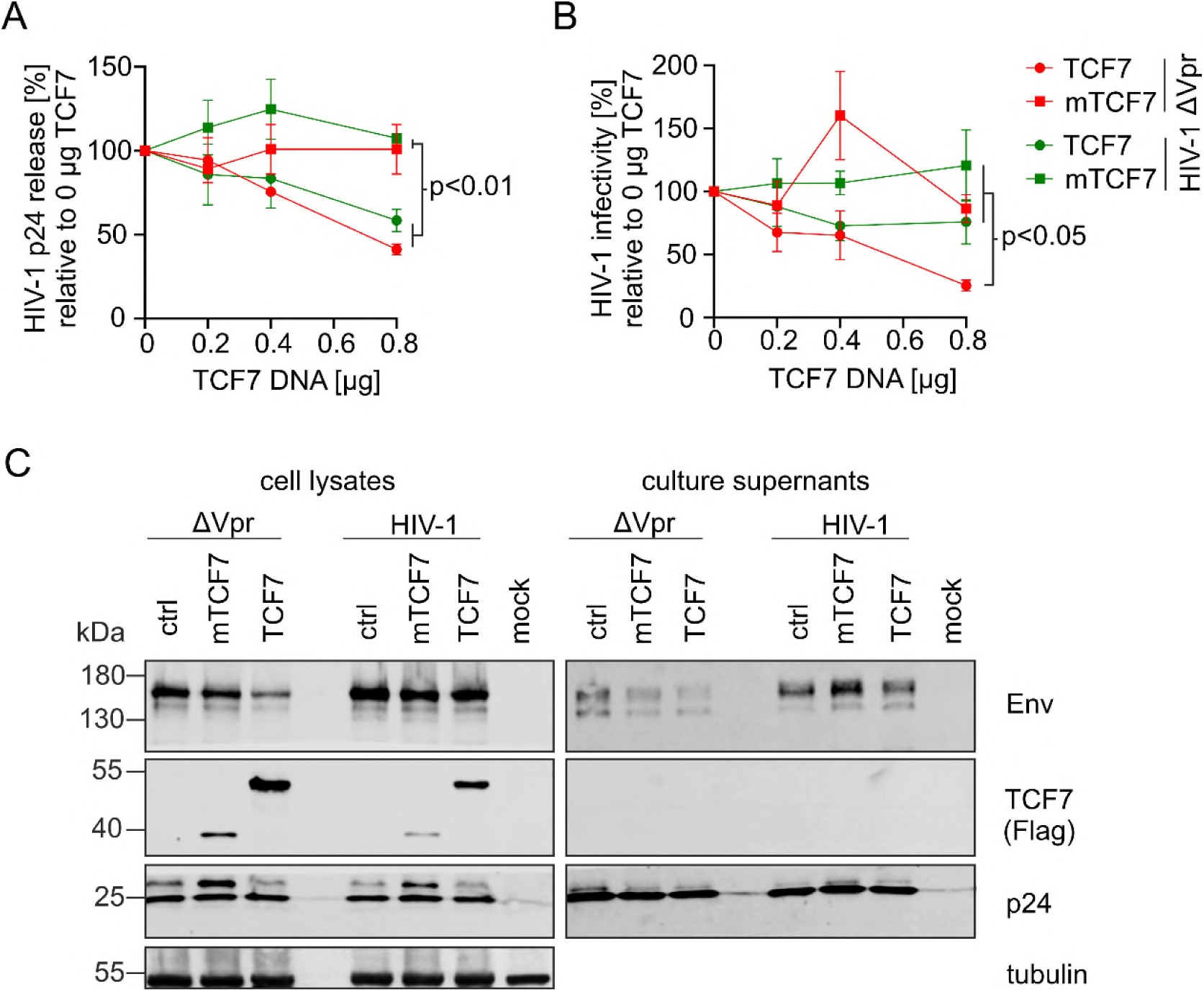
TCF7 modulates HIV-1 particle release and viral infectivity. (A) Expression plasmids encoding full-length TCF7 or truncated TCF7 (mTCF7), together with HIV-1 proviral constructs encoding functional *vpr* or defective *vpr*, were transfected into HEK293T cells. TCF7 refers to the long TCF7 isoform2 which contains the β-catenin-binding site, whereas mTCF7 refers to the truncated TCF7 variant. At 48 h post-transfection, virus-containing supernatants were harvested and quantified by p24 ELISA. HIV-1 particle release was normalized to the corresponding condition transfected without TCF7 (0 µg). Data are shown as the mean ± SD of n=3 biological replicates. Statistical significance was determined using a one-way ANOVA with Fisher’s LSD test.(B) Viral stocks generated in (A) were used to infect LC5-RIC reporter cells. Infection was quantified as the percentage of dsRed+ cells. Relative HIV-1 infectivity was calculated by normalization to virus stocks produced in the absence of TCF7 (0 µg). Data are shown as the mean ± SD of n = 4 (HIV-1) and 6 (ΔVpr) biological replicates. Statistical significance was determined using one-way ANOVA with Fisher’s LSD test. (C) Western blot analysis of virus- producing cells and virus stocks collected at 48 hpt from the condition transfected with 0.8 µg TCF7. Env, FLAG-tagged TCF7, and HIV-1 p24 were detected. Tubulin was used as a loading control for the virus-producing cell lysates.

### CD4^+^ T cell differentiation state determines HIV-1 permissiveness and Vpr promotes loss of stem-like memory CD4^+^ T cells

Given TCF7’s role in promoting stemness and preventing effector differentiation [47], we next assessed whether Vpr-induced TCF7 degradation influences the distribution of T cell subsets. Thereby, we analyzed six recognized T cell subsets: Naïve T cells (T_N_) have not yet encountered antigen and retain a high degree of lineage plasticity, whereas stem cell memory T cells (T_SCM_), despite being antigen-experienced, maintain their capacity for self-renewal [48, 49]. Central memory T cells (T_CM_), having already encountered antigen, respond rapidly upon re-exposure to antigen [50, 51]. Effector memory (T_EM_) and terminal effector (T_EFF_) T cells are more differentiated, have lost the lymph node homing receptor CD62L, and are characterized by immediate cytotoxic response. As a result, T_EM_ cells differentiate into T_EFF_ cells under sustained antigen exposure, with the latter being prone to exhaustion [49, 52]. Finally, terminally differentiated effector memory RA+ T cells (T_EMRA_) represent an exhausted, end-stage effector population [53]. To this end, primary CD4⁺ T cells were stimulated with IL-7 and IL-15, conditions that render cells permissive to HIV-1 infection without triggering TCR-mediated activation [54, 55]. Cells were infected with mock, HIV- 1, or ΔVpr , followed by analysis of TCF7 expression, p24 expression, and pairing of surface molecules like CD95, CD45RA, and CD62L to define distinct subsets at 2 to 7 days post-infection (Supplementary Figure S2) [49]. Two days post-infection was chosen to capture early effects, while phenotypic changes were expected from 5 dpi onward [55]. We first analyzed for overall effects in the cultures, without pre-gating for HIV-1 infected cells based on Gag-staining (Fig. 5A). Analysis of TCF7 expression revealed that T_SCM_ cells displayed the highest TCF7 levels, followed by T_CM_ and T_N_, with progressively lower expression in T_EM_ and T_EFF_ cells. The lowest TCF7 expression was observed within the T_EMRA_ population (Figure 5B). We next compared the overall infection for the CD4^+^ T cell subsets. HIV-1 permissiveness, as determined by infection levels at 2 dpi, mirrored this gradient in reverse: infection levels followed the order T_N_ < T_EMRA_ < T_SCM_ < T_CM_ < T_EM_ < T_EFF_, indicating that the more differentiated subsets are more susceptible to HIV-1 (Fig. 5C). A similar pattern was observed in the absence of Vpr; however, ΔVpr-infected cells showed comparable permissiveness for HIV-1 in T_SCM_ and T_CM_ subsets (Fig 5D). Next, we hypothesized that HIV-1 not only shows different permissiveness across subsets, but also that infection itself alters the phenotype of the T cell population. The amount of CD95^+^ cells over time stayed high, with a slight reduction upon 2dpi and 7dpi that was more pronounced in ΔVpr-infected cells (Supplementary Fig. S3A). CD62L was, as expected, significantly downregulated upon HIV-1 infection, independent of Vpr, with more robust downregulation present from 5dpi on (Supplementary Fig. 3B). In contrast, CD45RA expressing cells increased upon infection (Supplementary Fig. S3C). To address whether HIV-1 infection influences T cell differentiation and whether Vpr may play a role in directing T cell differentiation, subset distributions were examined. Under mock conditions, the majority of cells fell, as expected, within the T_CM_ and T_N_ subsets (Fig. 5E) [54, 56] and differentiated toward T_SCM_ from 2 to 7 days of culturing. Following HIV-1 infection, independent of Vpr, the distribution at 2 dpi resembled that of mock conditions, with high proportions of T_N_, T_SCM_, and T_CM_ cells. By 5 dpi, however, HIV-1-infected cultures displayed a marked shift; in detail, T_N_, T_SCM_ and T_CM_ proportions strongly decreased, and T_EM_, T_EFF_, and T_EMRA_ populations were enriched compared to the mock condition (Fig. 5E, middle panel). In ΔVpr-infected cells we observed a similar shift in subset frequencies, which was generally less pronounced as compared to HIV-1 infection and resulted especially at 5 dpi in increased T_SCM_ and reduced T_EFF_ subsets (Fig. 5E, central vs lower panel). Together, the data indicate that the differentiation states of CD4+ T cells strongly influence HIV-1 permissiveness. Of note, HIV-1 infection itself reshapes the composition of the T cell pool, with Vpr contributing to the accumulation of differentiated and hence HIV-1 permissive subsets, such as T_EM_ and T_EFF_, while T_SCM_ seem depleted.

**Figure 5.**
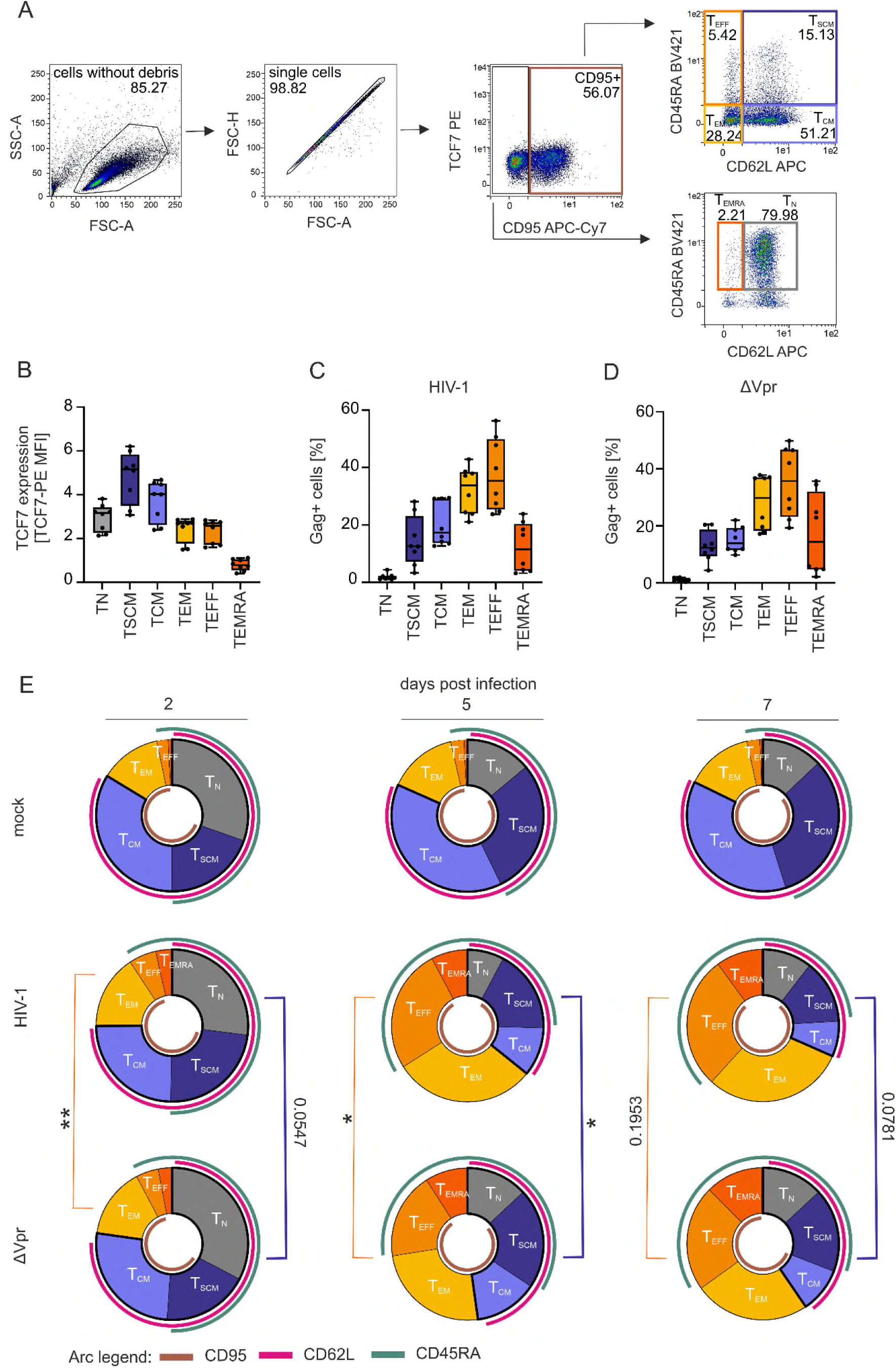
TCF7 expression and HIV-1 susceptibility across primary CD4^+^ T-cell subsets. Primary CD4^+^ T cells were maintained with IL-7 and IL-15 (20 ng/mL each), infected with HIV-1 or ΔVpr, and harvested between 2- and 7-days post-infection (dpi.) (A) Gating strategy used to identify the different CD4^+^ T-cell subsets according to their combined expression of CD95, CD62L, and CD45RA. (B) TCF7 expression, presented as median fluorescence intensity (MFI), within each individual T-cell subset in mock-infected cells harvested at 2 dpi. Box plots show the minimum to maximum values. n=8. (C and D) Susceptibility of each individual T-cell subset to HIV-1 infection, determined as the percentage of Gag+ cells at 2 dpi following infection with (C) HIV-1 or (D) ΔVpr. Box plots show the minimum to maximum values. n=8. (E) Distribution of the different T-cell subsets under mock-infected, HIV-1-infected, and ΔVpr-infected conditions at 2, 5, and 7 dpi. T-cell subset distributions were visualized as pie charts, with arcs indicating CD95^+^, CD62L^+^, and CD45RA^+^ populations, based on mean values from eight independent biological replicates. To assess differences in subset composition between HIV-1 and ΔVpr infection, we compared less differentiated (T_N_, T_SCM_, T_CM_) and more differentiated (T_EM_, T_EFF_, T_EMRA_) T cell populations using a Wilcoxon matched-pairs signed rank test. Comparisons between less differentiated subsets are highlighted in blue, and those between differentiated subsets in orange. Statistical significance is indicated as *p < 0.05 and **p < 0.01.

### TCF7 depletion recapitulates Vpr-dependent loss of T_SCM_ cells

Next, we asked whether TCF7 depletion in the context of HIV-1 ΔVpr infection, where TCF7 protein levels remain high, would recapitulate the Vpr-dependent phenotype of T_EM_ and T_EFF_ accumulation and T_SCM_ depletion. Since changes in T cell subsets were most apparent at later time points, the analysis focused on day 5 post-infection. Primary CD4^+^ T cells were cultured in IL-7 and IL-15 and treated with either non-targeting LNA or TCF7-targeting LNA 6 days prior to infection with HIV-1 or ΔVpr, followed by flow cytometry analysis to determine T cell subset distribution (Fig. 6A). LNAs are short DNA antisense oligonucleotides, so-called Gapmers, with locked nucleic acids flanking the target sequence of interest, that are taken up by T cells via gymnotic uptake [57]. Efficient TCF7 knockdown was confirmed in mock-treated and HIV-1 ΔVpr- infected cells (Fig. 6B and Supplementary Fig. S4A and B). As expected, HIV-1 infection resulted in the strongest reduction in TCF7 expression, leaving only approximately 20% of cells TCF7- positive at 5 dpi. TCF7 knockdown correlated with a reduction in CD95-expressing cells, independent of infection (Supplementary Fig. S4C), it did not influence CD62L expression (Supplementary Fig. S4D) and decreased the amount of CD45RA-positive cells in the ΔVpr condition (Supplementary Fig S4E). In non-infected (mock-treated) cultures, TCF7 knockdown increased the frequency of phenotypic T_N_ cells while decreasing the frequency of T_SCM_ (Fig. 6C). This is in line with the concept that TCF7 is required for the transition of naïve cells toward a T_SCM_-like state [58]. Upon HIV-1 infection, TCF7 knockdown did not alter the overall distribution of CD4^+^ T cell subsets. In contrast, in ΔVpr-infected cultures, where TCF7 expression is maintained, LNA-mediated knockdown of TCF7 reduced the proportion of T_SCM_ cells (Fig. 6C, right panel). TCF7 knockdown also increased p24 release into the culture supernatant in ΔVpr-infected cultures, as measured by p24 ELISA (Fig. 6D). This is consistent with our previous observations (Fig. 4A) and the results support the idea that loss of TCF7 promotes viral particle production, whereas preserved TCF7 expression reduces efficient virus release. When analyzing infected (Gag+) and non-infected bystander cells (Gag-) separately (Fig. 6E-J and Supplementary Fig. S5), TCF7 knockdown was associated with increased T_N_ and T_EMRA_ frequencies in both population (Fig. 6E and J), similar to the bulk culture (Fig. 6C). T_N_ frequency was low within the Gag+ population as compared to the Gag- population (Fig. 6E, left; Supplementary Fig. S5A). Within the T_SCM_ compartment, a clear Vpr- and TCF7-dependent pattern emerged (Fig. 6F). T_SCM_ frequencies were lower in HIV-1 infected cells than in ΔVpr-infections. For HIV-1, additional TCF7 knockdown did not further reduce T_SCM_ frequencies. By contrast, in ΔVpr-Gag+ infected T cells, where TCF7 expression is preserved, TCF7 knockdown reduced the T_SCM_ compartment to levels observed in HIV-1 infections (Fig 6F, left). A similar pattern was observed in the bystander population, with TCF7 KD decreasing the T_SCM_ compartment in HIV-1 and ΔVpr-infected cultures (Fig. 6F, right). These data indicate that TCF7 is required to maintain T_SCM_-like features and suggest that Vpr-mediated TCF7 degradation contributes to the loss of this population during HIV-1 infection. While T_CM_ were largely unaffected in this analysis (Fig. 6G), T_EM_ frequencies were enriched in HIV-1 infection, reaching statistical significance in the bystander population, irrespective of TCF7 knockdown (Fig. 6H). Moreover, in Gag+ cells, HIV-1 infection showed a trend to higher T_EFF_ frequencies than ΔVpr infection (Fig. 6I, left). Similarly, in the bystander population, ΔVpr-infected cultures contained fewer T_EFF_ cells than HIV-1-infected cultures and the frequency of T_EFF_ was increased upon TCF7 knockdown (Fig 6I, right). Together, our findings suggest that TCF7 restrains effector differentiation and that Vpr-dependent loss of TCF7 contributes to a shift away from a stem-memory-like phenotype toward a more differentiated effector state.

**Figure 6.**
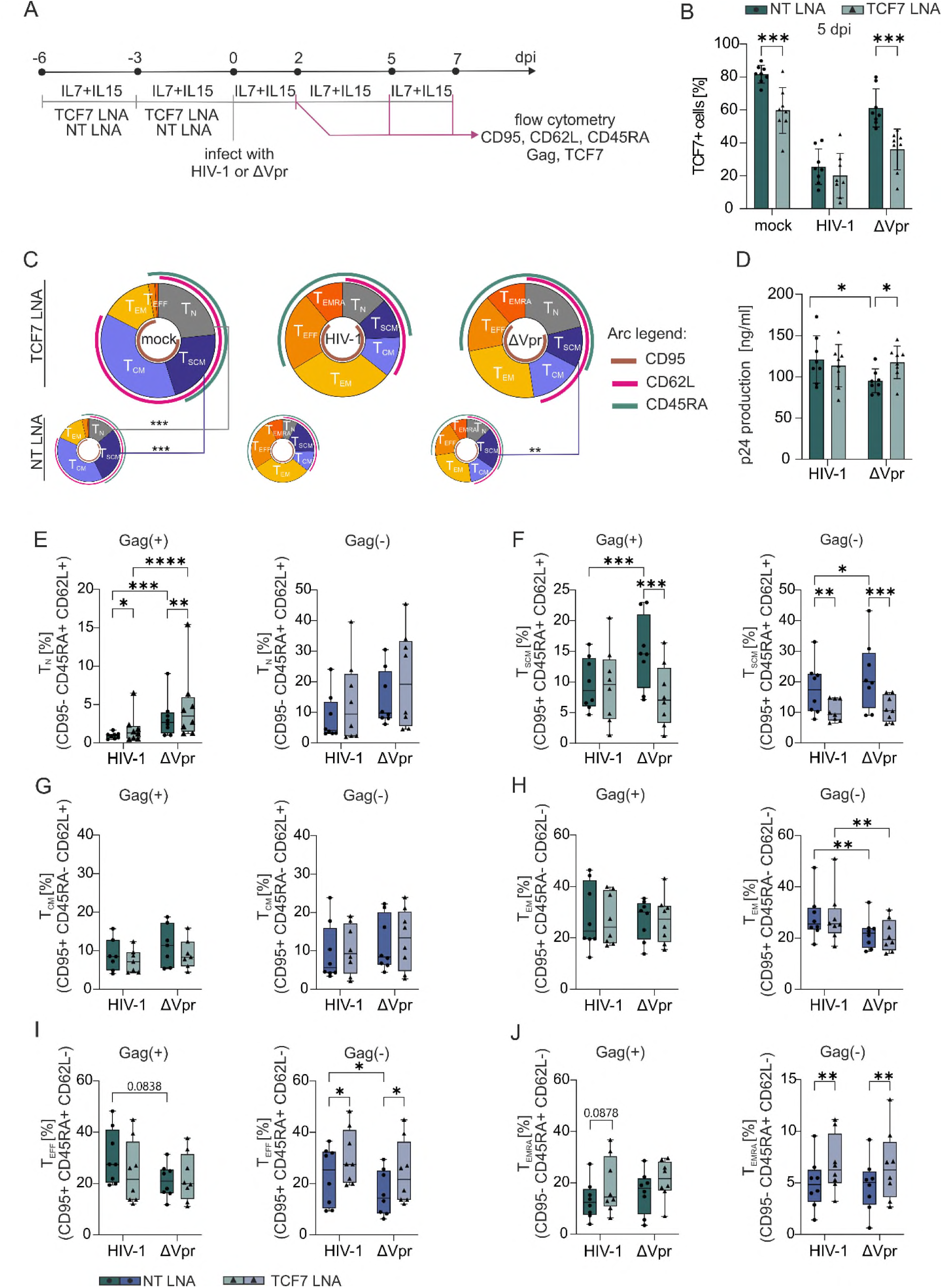
TCF7 knockdown alters CD4^+^ T-cell subset distribution. (A) Schematic representation of the experimental setup. Primary CD4^+^ T cells were stimulated with IL-7 and IL-15 (20 ng/mL each) and treated for 6 days with 6 µM non-targeting (NT) Gapmers containing flanking locked nucleic acids (LNAs) or with 3 µM each of two TCF7-targeting Gapmers containing flanking LNAs. Gapmers are hereafter referred to as LNAs. On day 3, the medium was replaced with fresh medium supplemented with IL-7 and IL-15, and the LNAs were readministered. Cells were infected with HIV-1 or ΔVpr and harvested at 2-, 5-, and 7-days post-infection (dpi) for flow-cytometric assessment of CD95, CD62L, CD45RA, TCF7, and HIV-1 p24 expression. Cells received fresh medium supplemented with IL-7 and IL-15 every 2–3 days. (B) TCF7 knockdown efficiency was determined at 5 dpi by intracellular flow-cytometric staining of TCF7. n=8 (Mean ± SD). Statistical significance was determined using paired two-way ANOVA with Šídák’s multiple- comparison test. ***p < 0.001.(C) Pie charts showing the distribution of the T-cell subsets at 5 dpi. Larger pie charts represent the TCF7 LNA-treated samples, and smaller pie charts represent the NT LNA-treated samples. Mean values from n=8 were used to visualize the T-cell subset distributions. Additional arcs indicate the expression of CD95, CD62L, and CD45RA. Statistical significance was determined using paired two-way ANOVA with Šídák’s multiple-comparison test. **p < 0.01; ***p < 0.001. (D) HIV-1 p24 release into the culture supernatant at 5 dpi was quantified by p24 ELISA. n=8 (Mean ± SD). Statistical significance was determined using a paired two-way ANOVA with Šídák’s multiple-comparison test. *p < 0.05. (E–J) Distribution of the indicated T-cell subsets within infected Gag (+) cells, shown in green, and bystander Gag (−) cells, shown in purple, comparing the effects of TCF7 knockdown using LNAs. (E) T_N_, (F) T_SCM_, (G) T_CM_, (H) T_EM_, (I) T_EFF_, and (J) T_EMRA_ cells are shown. Box plots extend from the minimum to the maximum (n=8). Statistical significance was determined using a paired two-way ANOVA with Šídák’s multiple- comparison test. *p < 0.05; **p < 0.01; ***p < 0.001; ****p < 0.0001.

Since Vpr-mediated TCF7 degradation was associated with loss of T_SCM_-like cells and enrichment of effector-like populations, we assessed cytokine secretion at 5 dpi, focusing on the effector-associated cytokines IFNγ and GM-CSF (Supplementary Fig. S6A). HIV-1-infected cultures secreted higher levels of IFNγ and GM-CSF than ΔVpr-infected cultures. In addition, TCF7 knockdown increased IFNγ and GM-CSF secretion in mock and ΔVpr conditions. These data further support a model in which Vpr-mediated loss of TCF7 promotes effector-like differentiation and function in HIV-1-infected CD4^+^ T cells.

### TCF7 depletion enhances activation responsiveness of CD4^+^ T cells

Vpr-mediated TCF7 degradation correlated with loss of T_SCM_-like characteristics, increased effector-like populations, and elevated secretion of effector-associated cytokines (Fig. 6 and S6), We therefore investigated whether TCF7 loss modulates the activation potential of CD4^+^ T cells. We quantified CD69 expression, an early activation marker induced by T cell stimulation, at 2 days post-infection to assess activation under IL-7 and IL-15 culture conditions. In a parallel approach, cells were stimulated with PHA three days before infection and three days after LNA-mediated TCF7 knockdown to determine whether TCF7 depletion affects activation responsiveness to a subsequent activating stimulus (Fig. 7A). The TCF7 knockdown under IL-7/IL-15 conditions was efficient (Fig. 7B). In contrast, LNA-mediated knockdown no longer resulted in a significant additional decrease in TCF7 levels upon PHA stimulation (Fig. 7C), potentially due to activation-associated downregulation of TCF7, thus masking the knockdown efficiency (Fig. 7C). As expected, HIV-1 infection increased the frequency of CD69-expressing cells (Fig. 7D; upper panel and Fig. 7E). A non-significant increase in CD69 compared to mock was observed in ΔVpr infections with no major differences detected in TCF7 knockdown conditions. By contrast, upon additional PHA stimulation, TCF7 knockdown significantly increased the frequency of CD69- expressing cells, particularly in mock and ΔVpr-infected cultures to levels comparable to HIV-1 infections (Fig. 7D, lower panel and Fig. 7F). Together, these findings suggest that TCF7 depletion enhances activation responsiveness of CD4^+^ T cells following TCR-stimulation. Thus, Vpr-mediated TCF7 degradation may lower the activation threshold of CD4^+^ T cells and promote the acquisition of an activated, effector-like phenotype during HIV-1 infection.

**Figure 7.**
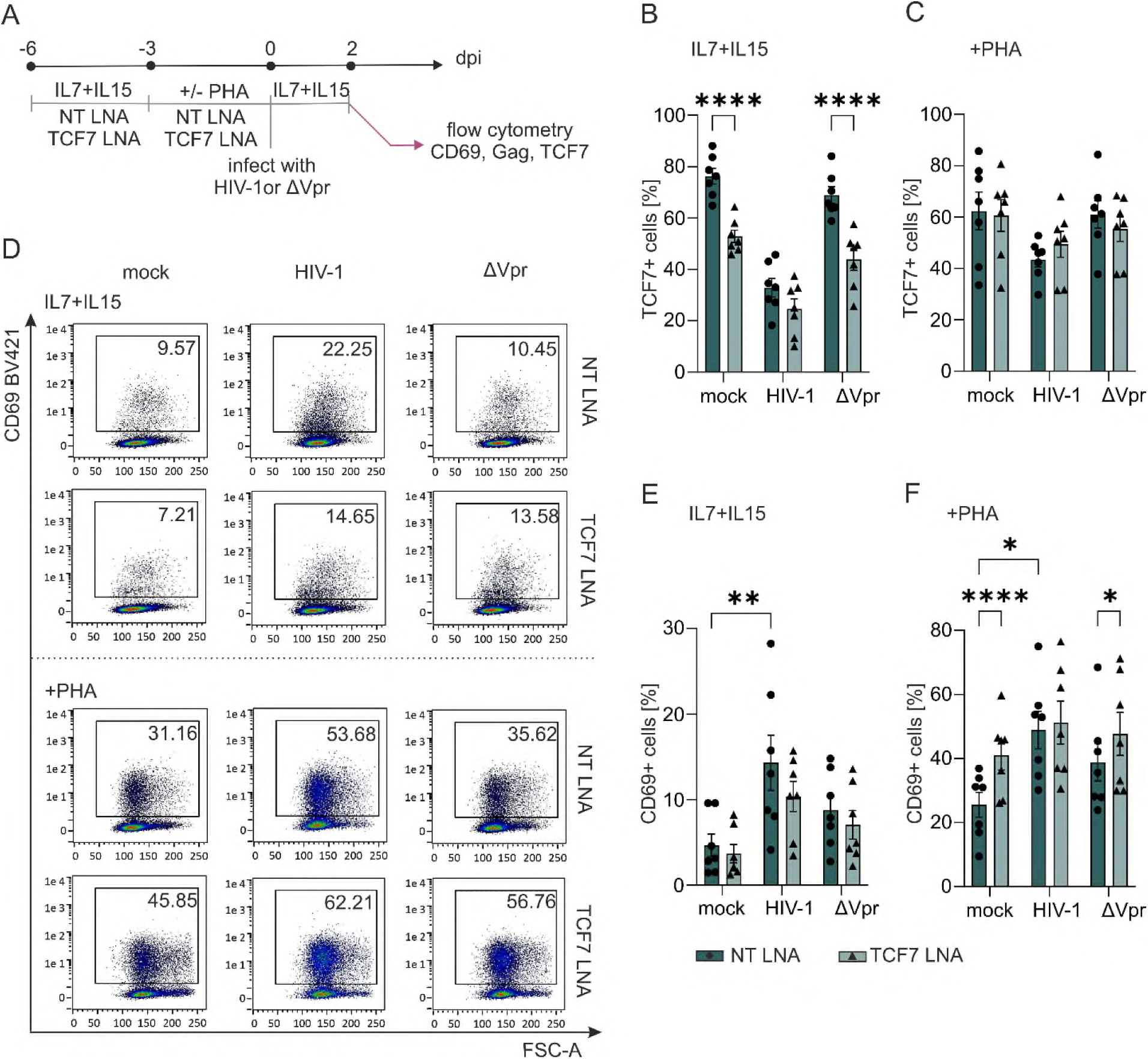
TCF7 knockdown modulates CD69 expression under cytokine- and mitogen- stimulating conditions. (A) Schematic representation of the experimental setup. Primary CD4^+^ T cells were stimulated with IL-7 and IL-15 (20 ng/mL each) and treated for 6 days with 6 µM non-targeting (NT) Gapmers containing flanking locked nucleic acids (LNAs) or with 3 µM each of two TCF7-targeting Gapmers containing flanking LNAs. On day 3, the medium was replaced with fresh medium supplemented with IL-7 and IL-15, and the LNAs were readministered. Cells were either maintained under IL-7 and IL-15 stimulation alone or additionally stimulated with PHA (1 µg/mL). Cells were infected with HIV-1 or ΔVpr and harvested at 2 dpi for flow-cytometric assessment of CD95, CD62L, CD45RA, TCF7, and HIV-1 p24 expression. (B and C) TCF7 knockdown efficiency was determined by intracellular flow-cytometric staining of TCF7 in cells maintained with (B) IL-7 and IL-15 alone or additionally stimulated with (C) PHA. n=7 (Mean ± SD). Statistical significance was determined using a paired two-way ANOVA with Šídák’s multiple-comparison test. ****p < 0.0001. (D) Representative flow-cytometry plots showing CD69 expression following infection and TCF7 knockdown under IL-7 and IL-15 stimulation alone (upper panel) or following additional PHA stimulation (lower panel). Quantification of CD69+ cells under (E) IL-7 and IL-15 stimulation alone or (F) following additional PHA stimulation. n=7 (Mean ± SD). Statistical significance was determined using a paired two-way ANOVA with Šídák’s multiple-comparison test. *p < 0.05; **p < 0.01; ****p < 0.0001.

## DISCUSSION

Vpr is known not only to drive striking transcriptional reprogramming in primary CD4^+^ T cells but also to modulate the host cell proteome through its interaction with the DCAF1 and CUL4A ubiquitin ligase complex, thereby targeting host proteins for proteasomal degradation [1, 6, 17, 42]. While Vpr’s role in supporting replication is less prominent in cell culture models but is becoming more relevant in primary HIV-1 target cells [2], we decided to investigate the protein landscape of primary CD4^+^ T cells and how it is remodeled upon HIV-1 infection. Here, we present a comprehensive proteomic approach to identify host cell factors degraded by Vpr. Confirming our proteomic approach, we found known HIV-1 targets, such as CD4 [33, 34], APOBEC3F [36], and PPP2 [35], being degraded upon HIV-1 infection. Comparing HIV-1 with intact or defective *vpr* open reading frame, we identify TCF7 as a novel Vpr target. We found that this Vpr function is conserved across HIV-1 subgroups and primary alleles. Vpr is packaged into HIV-1 virions and is one of the most abundant proteins within them, thus present during early infection events. In fact, Vpr alone, poised to the host cell via incoming virions, is in fact sufficient to reduce TCF7 protein levels, with TCF7 reduction in the full-length viral context already happening at 4 hours post infection (Figure 2E-G). Vpr-mediated modulation of TCF7 does not require de novo Vpr transcription and protein synthesis, allowing HIV-1 to directly and immediately modulate the cell, even in non-activated cells, which are not productively infected or non-infected bystander cells. Using different Vpr mutants to get a deeper insight into structure-function, we identified that Vpr-mediated TCF7 reduction was dependent on Vpr to interact with DCAF1, since Q65R and C76A are defective for TCF7 and fail to bind to DCAF1 [3, 39, 59–61]. In line with this, in HIV-1 infection, TCF7 levels were restored in the presence of MG132, while blocking the lysosomal pathway did not restore protein expression, and TCF7 mRNA levels were not decreased in primary CD4^+^ T cells upon HIV-1 infection. This suggested that TCF7 is a canonical Vpr target, which is directed via the DCAF1 and CUL4A ubiquitin ligase complex towards proteasomal degradation. However, we did not detect a direct TCF7 interaction with either Vpr or DCAF1. This suggests that, although TCF7 degradation is proteasome-mediated and DCAF1-dependent, TCF7 is neither a direct binding partner of DCAF1 nor Vpr. Hence, TCF7 might be targeted for proteasomal degradation via an indirect pathway, e.g., through Vpr’s role in activating ATM or ATR signaling pathways [62–64], with the latter recently shown to be the underlying cause of CCDC137 depletion accompanying nucleolar stress [65]. Of note, CCDC137 degradation was also restored upon MG132-mediated proteasome inhibition without direct DCAF1 interaction, similar to TCF7 [65]. Our current working models that will be experimentally explored are that Vpr degrades a TCF7 stabilizer via the DCAF E3-ubiquitin ligase machinery or induces DCAF1-coupled cellular responses that activate the endogenous TCF7 turnover machinery. In this context, ubiquitin-independent proteasomal degradation of TCF7 directed via the substrate recognition compontent of the E3 ubiquitin ligase complex pVHL [66, 67] as well as neddylation-dependent proteasomal degradation has been described [68].

But what is the functional importance of TCF7 degradation in HIV-1 infection? HIV-1 evolved multiple mechanisms to evade so-called host restriction factors that are often induced by interferon and exert direct inhibitory activity on multiple steps of the viral replication cycle [69, 70]. While TCF7 is not interferon-regulated (Interferome database [71]; https://isg.data.cvr.ac.uk/), our data from HIV-1 producing 293T cells that overexpress TCF7 (Fig. 4A) as well as from IL-7 and IL-15 activated CD4^+^ T cells (Fig. 6D) suggest that high levels of TCF7 expression directly translate into less viral particle (p24)-release. This comes alongside a reduced specific particle infectivity, as high levels of TCF7 seem to interfere with optimal Env expression and incorporation into viral particles (Fig. 4B and C). Of note, Vpr-regulated optimization of Env expression and hence increased HIV-1 spread has been described exclusively for macrophages thus far, via targeting the transcription factor PU.1 for degradation [72], suggesting that Vpr evolved a similar mechanism in T cells via TCF7. Nevertheless, we do not consider TCF7 a classical restriction factor but favor a model in which Vpr degrades TCF7 to promote T cell differentiation into highly HIV-1 permissive subsets that fail to orchestrate an efficient antiviral immune response (Fig. 8). Accordingly, we found that more differentiated cells, such as effector memory and effector cells, seem more permissive to HIV-1 infection, whereas naive and T_SCM_ cells are less susceptible (Fig. 5C and D), consistent with previous reports [55, 73]. Intriguingly, TCF7 expression followed the inverted pattern, being most abundantly expressed in T_SCM_ and lowest in differentiated subsets T_EM_, T_EFF_, and T_EMRA_ cells (Fig. 5B). This is in line with TCF7 mainly described as a transcription factor being important for stemness and a negative regulator of effector cell responses in CD4^+^ T cells [37].

**Figure 8.**
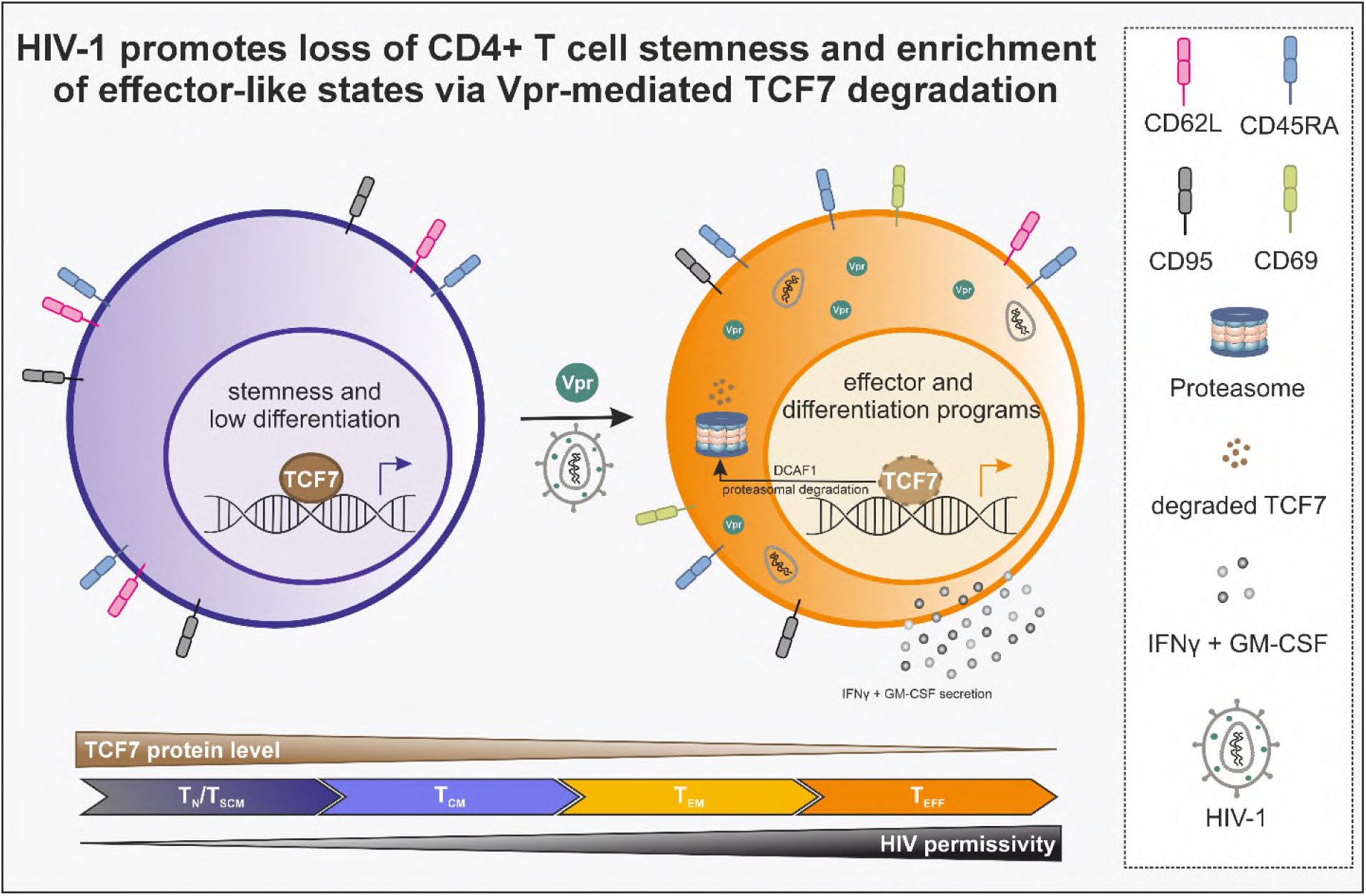
Graphical Abstract: HIV-1 Vpr-mediated loss of CD4^+^ T cell stemness via TCF7 degradation. Vpr targets TCF7 for proteasomal degradation via a DCAF-dependent non- canonical pathway, resulting in a loss of CD4^+^ T cells’ stemness and enrichment of effector-like states, associated with increased IFNg and GM-CSF secretion. Furthermore, TCF7 protein expression inversely correlated with HIV susceptibility, indicating that differentiated effector-like states are more susceptible to HIV-1 infection. Figure was generated using CorelDraw.

By combining subset-defining markers, we found that HIV-1 infection induced a shift toward an effector phenotype with less T_N_ and T_SCM_ and corresponding increase in T_EM_ and T_EFF_ subsets in both bystander and infected cells (Fig. 5E and Fig. 6). This phenotypic shift is mediated by degradation of TCF7 upon infection. In support of this, the proportion of T_SCM_ was markedly increased in HIV-1 ΔVpr-infections, and brought to HIV-1 levels by knocking-down TCF7 (Fig. 6F). HIV-1-induced acquisition of an effector-like phenotype and loss of T_SCM_, mediated by Vpr, was associated with increased secretion of IFNγ and GM-CSF. IFNγ is a canonical Th1/effector cytokine and has previously been shown to be negatively regulated by TCF7 expression [28], while elevated GM-CSF reflects enhanced inflammatory effector activation [74, 75]. Together, these findings indicate that HIV-1 triggers, via TCF7 reduction, exit from T_SCM_-like CD4^+^ T cell states towards differentiated and highly permissive T cell subsets. Indeed, T_SCM_ can differentiate into T_CM_ and T_EM_ following antigen encounter [48], and T_EM_ can subsequently differentiate into T_EFF_ [76]. Our results support a model, in which T_SCM_ cells are characterized by a less differentiated and functionally restrained state [77], and their activation responsiveness is enhanced via TCF7 depletion, mediated by Vpr (Fig. 7 and Fig. 8). Consequently, Vpr but not TCF7 downmodulation per se contributes to early activation-associated remodeling of primary CD4^+^ T cells, consistent with reported Vpr-dependent NFAT activation and CD69 induction [3, 4]. Thus, Vpr-mediated TCF7 degradation may cooperate with Vpr-induced activation pathways to lower the threshold for effector-associated CD4^+^ T cell responses during HIV-1 infection. Of note, in PLWH, TCF7 expression was lower in rapid progressors as compared to non-progressors and healthy controls [78]. On top, TCF7 levels correlated negatively with viral loads and CD4^+^ T cell counts, with low TCF7 expression predominantly in T_EM_ and T_EFF_. While TCF7 mRNA appeared also reduced, the latter study provides the *in vivo* framework reflecting our main findings [78]. In the context of HIV- 1 pathogenesis, TCF7 may help to preserve a renewable and functionally competent CD4^+^ T cell pool, and its Vpr-mediated loss, in infected as well as non-infected bystander CD4^+^ T cells, could induce a pre-depletion imposed progressive loss of HIV-specific helper CD4^+^ T cell function and T cell exhaustion, also in CD8^+^ T cells [79]. Evidence for this hypothesis comes from LCMV mouse models showing that TCF7-expressing CD4^+^ T cells, depending on BCL6, are progenitors needed to maintain CD4^+^ effector and T helper cell responses, as well as T follicular helper cells [80, 81]. Conversely, a setting that preserves T cells with high TCF7 levels and T_SCM_-like properties might favor HIV-1 persistence and latency [82, 83].

Overall, our findings support a model in which TCF7 is required to maintain or establish the T_SCM_- like state. In the absence of Vpr, TCF7 expression is preserved, and T_SCM_ cells are maintained more efficiently. Progressive infection is linked to the disruption or loss of less-differentiated memory compartments, whereas preservation of T_SCM_ and T_CM_ compartments is associated with better CD4^+^ T-cell homeostasis in viremic non-progressors [84]. Thus, Vpr-mediated TCF7 downregulation may represent a mechanism by which HIV-1 remodels the CD4^+^ T-cell differentiation hierarchy, reducing stem-memory features while favoring effector-like states. Thereby, T cells are shifted away from a quiescent, stem-memory-like state toward a more HIV-1 permissive effector-like phenotype (Fig. 8). Furthermore, loss of CD4^+^ T-cell stemness contributes to immune dysfunction and impaired antiviral immune response.

## MATERIAL & METHODS

### Proviral constructs and plasmids

HIV-1 constructs based on the pBR-NL4.3 backbone, including variants encoding intact or deleted *vpr* open reading frames, were used as previously described [3, 85, 86]. Vpr point mutants were expressed in a pCG-IRES-GFP vector as previously described. NL4.3 Vpr Q65R was generated by cloning a synthesized Vpr Q65R with flanking MluI and XbaI restriction sites sequence (Eurofins) into the pCG-IRES-GFP backbone using MluI and XbaI restriction sites. NL4.3 Vpr R85Q was generated by PCR-based site-directed mutagenesis (fw_Vpr_XbaI: gcgTCTAGA atg gaa caa gcc cca gaa gac caa ggg cc; rev_VprR85QMIui: tgcACGCGT cta gga tct act ggc tcc att tct tgc tct cct ctg ctg agt aac gcc tat tc) incorporating overlapping MluI and XbaI restriction sites. All constructs were verified by Sanger sequencing. Vpr sequences from primary isolates cloned into a pCG-AU1-IRES-GFP backbone were kindly provided by Frank Kirchhoff (Ulm, Germany)[87]. GFP-expressing pWPI lentiviral vectors encoding a panel of primary HIV-1 Vpr sequences, along with cognate controls, were kindly provided by Éric A. Cohen (Montreal, Canada) [88]. Packaging plasmid psPAX2 (Addgene #12260) and VSV-G envelope plasmid pMD2G (Addgene #12259) were obtained from Addgene. Alternatively, pHIT-G expressing the vesicular stomatitis virus G protein (VSV-G) was used [89]. pcDNA3-HA-TCF1 (Addgene #40620) was kindly provided by Daniel Sauter (Tübingen, Germany) [90]. pcDNA3.1, pcDNA3.1+/C-(K)DYK_TCF7_FLAG and pcDNA3.1+/C-(K)DYK_TCF7v1_FLAG were ordered from GenScript. pCG-nef3* IRES GFP was described elsewhere [91].

### Cell lines

HEK293T cells were maintained in DMEM GlutaMAX (Gibco, Life Technologies) supplemented with 10% fetal calf serum (FCS; Gibco) and 1% penicillin and streptomycin (P/S; Life Technologies) at 37°C in a humidified 5% CO₂ atmosphere. LC5-RIC, indicating RIC as red- infected cells and LC5 as the parental cell line stably expressing CD4 and CXCR4 receptors and DsRed1 reporter, which is activated by HIV-1 Tat and Rev [46], was cultured in DMEM supplemented with 10% FCS and 1% P/S and additionally maintained under selection pressure by the addition of 0.74 mg/ml Gentamicin (G418; Thermo Fisher) and 0.125 mg/ml Hygromycin B (Applichem) at every second passage. SupT1 cells were cultured in RPMI-1640 GlutMAX supplemented with 10% FCS and 1% P/S under the same conditions.

### Primary CD4^+^ T cell isolation and culture

Primary CD4^+^ T cells were isolated from buffy coats obtained from anonymous healthy donors (ZKT Tübingen gGmbH, Tübingen, Germany). Donors provided written informed consent, and the use of blood-derived cells for research purposes was approved by the ethics committee of the University Hospital Tübingen (IRB #860/2023BO2). CD4+ T cells were enriched by negative selection using the RosetteSep Human CD4^+^ T Cell Enrichment Cocktail (StemCell Technologies) followed by density gradient centrifugation over Ficoll-Paque Plus (VWR). Isolated CD4^+^ T cells were maintained in RPMI-1640 GlutaMax supplemented with 10% FCS and 1% P/S. To maintain viability without activation, purified CD4^+^ T cells were cultured in the presence of 10 ng/ml IL-2 (StemCell Technologies) or, alternatively, 20 ng/ml IL-7 (Immunotools) and 20 ng/ml IL-15 (Immunotools). Where activated CD4^+^ T cells were required, cells were either pre-stimulated with 1 µg/mL phytohemagglutinin (PHA; Thermo Fisher Scientific) for 3 days before infection or stimulated over the TCR with 2 µg/mL plate-bound anti-CD3 (clone HIT3a; Biolegend) plus 1 µg/mL soluble anti-CD28 (clone CD28.2; Biolegend) in the presence of 10 ng/ml IL-2 3 days prior to infection.

### Viral stock production

Virus stocks were produced by transient transfection of HEK293T cells with NL4.3 provirus containing an intact or defective *vpr* ORF as described before [3, 86, 92]. Transfection was performed using either JetPrime (Polyplus) or polyethylenimine (PEI; Polysciences) in OptiMEM (Thermo Fisher), followed by a medium change at 4- or 16-hours post-transfection, respectively. To generate HIV-1 stocks with Vpr trans-complementation, *Vpr*- defective NL4-3 backbones were co-transfected with the appropriate Vpr expression plasmid (pCG-IRES-GFP or pCG-AU1-IRES-GFP) at a 4:1 ratio. VSV-G pseudotyped particles were produced by co-transfection of the proviral vector, Vpr expression plasmid where applicable, and pHIT-G (VSV-G) at a 10:1 ratio. For virus stocks produced in the presence of TCF7 overexpression, IRES-GFP proviral constructs with or without *vpr* were co-transfected with varying amounts of pcDNA3.1+/C-(K)DYK_TCF7_FLAG or pcDNA3.1+/C-(K)DYK_TCF7v1_FLAG, as indicated in the respective experiments. Empty pcDNA3.1 vector was used to equalize total DNA amounts across conditions.

Viral stocks were harvested at 24 hours post-transfection for VSV-G pseudotyped samples and at 48 hours post-transfection for all other samples. Vpr-expressing VLPs were generated using second-generation lentiviral packaging as previously described [7, 86, 88]. All viral supernatants were clarified by centrifugation at 3,200 × g for 10 minutes at 4°C to remove residual cells and debris. For infection of primary CD4^+^ T cells, HIV-1 NL4.3 stocks were concentrated by centrifugation through a 20% (w/v) sucrose cushion at 20,000 × g for 90 minutes to 2 hours at 4°C. Viral pellets were resuspended in FCS-free RPMI-1640 GlutMAX at 1/100 of the original volume, and viral stock concentration was assessed by HIV-1 p24 capsid ELISA as described below.

### Infection assays

For infection of SupT1 cells, 2 × 10⁶ cells/ml were seeded and infected directly with VSV-G-pseudotyped HIV-1 stocks encoding Vpr variants from different subtypes or with Vpr-expressing VLPs. Cells were harvested 24–48 hours post-infection for downstream analysis. For infection of primary CD4^+^ T cells, cells were seeded at 2–3 × 10⁶ cells/ml and infected with 200– 300 ng p24/ml, using equivalent p24 inputs for HIV-1 and *Δ*Vpr. Infection was enhanced by spinoculation at 800 × g for 45 minutes to 2 hours at 32°C. Cultures were washed 6- 16 hours post-infection and resuspended in RPMI-1640 with FCS and P/S supplemented with 10 ng/ml IL- 2 or 20 ng/ml IL-7 and 20 ng/ml IL-15. When infected cells were kept in culture for more than 2 days post-infection, the medium was changed every 2 to 3 days. Where indicated, cultures were treated with 100 nM Efavirenz (Sigma) or 250 nM Raltegravir (Sigma); 1 µM MG132 (AdipoGen Life Sciences), or 500 nM Bafilomycin A1 (AdipoGen Life Sciences), added 2 hours post-infection, with DMSO (Sigma) as vehicle control. Of note, Infection rates were either assessed using Western Blotting or intracellular anti-Gag staining and flow cytometry. For infection of LC5-RIC cells, cells were seeded on day prior to infection and infected with HIV-1 and ΔVpr viral stocks that were produced in the presence of varying amounts of TCF7. Infection was measured as DsRed1 reporter expression using Incucyte (with S3/SX1 G/R Optical Module, Sartorius). The amount of red expressing cells and confluency was quantified using Incucyte Base Analysis Software (Live-Cell Imaging and Analysis Software, Sartorius).

### p24 antigen quantification

To quantify HIV-1 p24 capsid antigen levels, virus-containing samples were inactivated by incubation with PBS containing 10% Triton X-100 (Merck Millipore) for 1 hour at 37°C. p24 concentration was measured using an in-house-developed p24 ELISA [93]. The absorbance was read using a Berthold TriStar2 S LB 942 multimode reader (Berthold Technologies GmbH & Co. KG). Of note, for quantification of p24 release into the supernatant of infected primary T cells in the context of TCF7 knockdown experiments, supernatants were collected at the day of cell harvest, spun down at 3,200 xg for 10 min at 4 °C to remove any debris, and frozen at -80 °C until further use.

### LNA-mediated knockdown

For TCF7 knockdown, Gapmer oligonucleotides with flanking locked nucleic acids at both ends targeting *TCF7* mRNA or a non-targeting control (NT) were designed. LNAs were ordered and HPLC-purified from Biomers. The sequences were as follows:

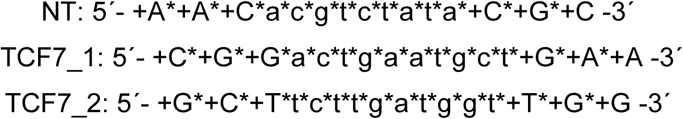

Asterisks indicate phosphorothioate backbone modifications, and plus signs indicate locked nucleic acid-modified bases. Primary CD4^+^ T cells in 20 ng/ml IL-7 and 20 ng/ml IL-15 were seeded at 1.5 × 10⁶ cells/ml. TCF7 expression was silenced using the combination of LNAs TCF7_1 and TCF7_2 (each 3 µM; 6 µM total), alongside an NT LNA control ( 6 µM). LNAs were added directly to the culture medium without a transfection reagent, as LNAs are taken up via gymnotic delivery. At day 3 post-treatment, cultures received a medium change, and LNAs were re-administered at the same concentrations. Where indicated, cells were additionally stimulated with 1 µg/ml PHA (Sigma) at this point. On day 6 post-initial LNA treatment, cells were distributed according to experimental conditions and infected with NL4.3 encoding intact or deleted *vpr,* as described above. Knockdown efficiency was assessed following infection at 2-7 dpi by flow cytometry.

### Flow cytometry

Infected primary CD4^+^ T cells were stained for surface expression of memory and activation markers essentially as before [92]. Cells were first blocked in PBS containing 10% FCS for 30 minutes at room temperature, then incubated with antibody cocktails diluted in FACS buffer (PBS containing 1% FCS) for 30 minutes at 4°C. The following anti-human antibodies were used for surface staining: CD95-APC/Cy7 (clone BX2; 1:50), CD62L-APC (clone DREG-56; 1:50), CD45RA-Brilliant Violet 421 (clone HI100; 1:50), and CD69-Brilliant Violet 421 (clone FN50; 1:50), all from BioLegend. Cells were washed thoroughly to remove unbound antibody. For intracellular detection of TCF7 and HIV-1 gag, cells were fixed and permeabilized using the eBioscience Fixation/Permeabilization Kit (Thermo Fisher Scientific) according to the manufacturer’s instructions, then stained with anti-TCF7-PE (clone 7F11A10; 1:50; BioLegend) and KC57-FITC (anti-Gag; 1:100; Beckman Coulter) in 1X PERM buffer supplemented with 10% FCS for 15 minutes on ice followed by 45 minutes at room temperature. All samples were acquired on a MACSQuant VYB or MACSQuant M10 (Miltenyi Biotec). Data was analyzed using FlowLogic software.

### LEGENDplex (Cytokine Release Assay)

Supernatant of HIV-1-infected primary CD4 T cells, maintained in RPMI supplemented with 20 ng/ml IL-7 and IL-15, treated with non-targeting LNA or TCF7-targeting LNAs, were collected at 5 dpi, spun down at 3,200 × g at 4 °C, and stored at - 80 °C until further use. In parallel, cells were harvested at the same time point, stained for different cell surface markers, and cell count was assessed. For cytokine quantification, undiluted supernatants were thawed on ice and analyzed using the LEGENDplex Human Anti-Virus Response Panel 1 Vo2 (Biolegend) according to the manufacturer’s protocol. Beads specific for interferons (α, β, γ, λ1, and α2), interleukins (1β, 6, 8, 10, and 12), TNF-α, IP-10, and GM-CSF were added to each sample in assay buffer. Samples were incubated for 2 hours at RT under shaking (800 rpm), washed twice with LEGENDplex 1X Washing buffer. Beads were incubated with the detection antibody for 1 hour at RT, followed by addition of the SA-PE antibody for 30 min at RT under shaking (800 rpm). Of note, the kit-provided Standard Cocktail was used as recommended, and RPMI media with FCS and P/S supplemented with 20 ng/ml IL7 and IL15 was used as a media background control. Samples and controls were performed in duplicates. After antibody incubation, beads were washed twice and fixed with 2% PFA for 10 min at RT, then washed twice with 1X Washing buffer. All samples were acquired on a MACSQuant M10 (Miltenyi Biotec). Data was analyzed using FlowLogic software (v8.3; Inivai). A standard curve for each cytokine was generated by applying a 5-parameter asymmetric sigmoidal curve fit, when applicable, using GraphPad Prism (v10.1.1). Mean fluorescence intensity values were interpolated to determine cytokine concentrations. To account for differences in cell counts between infected and non-infected samples, cytokine concentrations were normalized to the number of viable cells at the time of supernatant collection, as determined by flow cytometry. Cytokine secretion was extrapolated to 100,000 cells using the formula: (interpolated cytokine concentration × 0.025 ml / cell count) × 100,000 cells. Only IFNγ and GM-CSF measurements were included in the analysis, as other cytokines were at or near background levels despite the use of undiluted supernatant.

### Western Blotting

To assess Env, TCF7, p24, and Vpr protein expression, total protein from SupT1, primary CD4^+^ T, and HEK293T cells was lysed in RIPA buffer (140 mM NaCl, 10 mM Tris-HCl, 1 mM EDTA, 0.5 mM EGTA, 0.1% (w/v) Na-deoxycholate, 0.1% (w/v) SDS, 1% (v/v) Triton X-100, pH 7.4) supplemented with protease inhibitor (SIGMAFAST Protease Inhibitor Cocktail; Sigma-Aldrich) and phosphatase inhibitor (PhosSTOP; Merck Millipore) for 1 hour on ice. The cell lysates were centrifuged at 20,000 x g for 10 min at 4 °C. Supernatants were then boiled with 6X SDS loading dye (0.5 M Tris, pH 6.8, 2% bromophenol blue, 30% glycerol, 10% SDS, 0.6 M DTT) at 95 °C for 10 min. To assess Env and p24 protein levels of viral stocks, stocks were cleared, subsequently lysed with Triton X-100 (final concentration of 1% (v/v) Triton in PBS) for 30 min at 37 °C, and boiled with 6X SDS loading dye at 95 °C for 10 min. Proteins were resolved by SDS-PAGE on 10–12% Tris-Glycine or Bis-Tris gels using either SDS (25 mM Tris, 192 mM glycine, 0.1% (w/v) SDS; pH 8.3) or MES running buffer (50 mM Tris, 50 mM MES, 0.1% SDS, 1 mM EDTA; pH 7.3), respectively, followed by transfer to 0.2 µm nitrocellulose membranes (VWR) by wet transfer at 4°C. Membranes were blocked in 5% (w/v) skimmed milk in TBS for 1 hour at room temperature, then incubated overnight at 4°C with primary antibodies diluted in blocking buffer supplemented with 0.1% Triton X-100. Membranes were subsequently probed with IRDye-conjugated secondary antibodies diluted in blocking buffer supplemented with 0.1% Triton X-100 for 1- 2 hours at room temperature. Blots were imaged on a LI-COR Odyssey infrared imaging system. Membranes were re-probed with anti-GAPDH, anti-β-actin, or anti-tubulin as a loading control. The following antibodies were used: rabbit anti-TCF7 (clone: C63D9; 1:1,000; Cell Signaling); rabbit anti-DCAF1 (VPRBP; clone:D5K5V; 1:1,000; Cell Signaling); rabbit anti-β-Catenin (clone: D10A8; 1:1,000; Cell Signaling); rabbit anti-HIV1 p24 (clone:P131; 1:1,000; abcam); mouse anti-FLAG (clone:M2; 1:1,000; Merck); rabbit HIV-1 Vpr (1–50) antiserum (1:800; NIH HIV Reagent Program, Division of AIDS, NIAID #11836, contributed by Dr. Jeffrey Kopp), mouse HIV-1 envelope (clone: 16H3; 1:1,000; NIH HIV Reagent Program, Division of AIDS; NIH:16H3 mAB from Drs. Barton F. Haynes and Hua-Xin Liao), rat anti-GAPDH (clone: W17079A; 1:1,000; BioLegend), rabbit anti-β-tubulin (1:1,1000; Invitrogen); mouse anti-actin (clone: AC-40; 1:1,000; Merck) and IRDye 680RD- or 800CW-conjugated anti-rabbit, anti-mouse and anti-rat IgG secondary antibodies (1:15,000; LI-COR).

### Co-immunoprecipitation

To investigate TCF7–Vpr/DCAF1 protein-protein interactions, HEK293T cells were co-transfected with pcDNA3-HA-TCF1 and Vpr or Vpr Q65R expression constructs (pCG-IRES-GFP backbone) at a 1:1 ratio using JetPrime (Polyplus) according to the manufacturer’s instructions. To ensure equal DNA amounts, pCG-nef3*-IRES GFP. Medium was changed at 6 hours post-transfection to medium containing 1 µM MG132 or DMSO vehicle control. Cells were harvested 24 hours post-transfection and lysed in co-immunoprecipitation lysis buffer (Co-IP lysis buffer: 50 mM HEPES pH 7.5, 150 mM NaCl, 0.3% Igepal CA-630, 1 mM DTT) supplemented with protease inhibitor (SIGMAFAST Protease Inhibitor Cocktail; Sigma-Aldrich) and benzonase (250U; Merck Millipore) for 30 minutes on ice. Lysates were cleared by centrifugation at 20,000 ×g for 10 minutes at 4 °C, and an aliquot was taken as input control and boiled with 6X SDS loading buffer (2% (w/v) bromophenol blue, 30% (v/v) glycerol, 10% (w/v) SDS, 0.6M DTT, fill up with 0.5 M Tris; pH 6.8) at 95 °C. Cleared lysates were incubated with 20 µl anti-HA magnetic beads (Miltenyi Biotec) in a total volume of 400 µl Co-IP lysis buffer with protease inhibitor for 2 hours at 4°C under continuous rotation. Bead-bound complexes were captured over µMACS columns (Miltenyi Biotec) pre-equilibrated with the Co-IP lysis buffer. Columns were washed twice with 500 µl and once with 250 µl Co-IP lysis buffer. Bound proteins were eluted using 1X SDS loading buffer pre-heated to 95 °C. Input and elution samples were resolved by SDS-PAGE (Bis-Tris Gels) and transferred to nitrocellulose as described above.

### qRT-PCR

Total RNA was extracted from pre-stimulated, infected primary CD4^+^ T cells at 48 hours post-infection using the RNeasy Mini Kit (Qiagen) and reverse transcribed using the QuantiTect Reverse Transcription Kit (QIAGEN). RT-qPCR was performed in technical triplicate using the Luna Universal qPCR Master Mix (New England Biolabs). Each reaction contained 5 ng cDNA and 6 pmol of each primer. *TCF7* mRNA levels were quantified using QuantiTect Assay Primer (Hs_TCF7_1_SG; QIAGEN). *GAPDH* was used as the reference housekeeping gene with previously described primers (forward: 5′-TGCACCACCAACTGCTTAGC-3′; reverse: 5′- GGCATGGACTGTGGTCATGAG-3′) [94]. Reactions were run on a LightCycler® 480 (Roche) with an initial denaturation at 95°C for 10 minutes, followed by 45 cycles of 95°C for 10 seconds, 55°C for 15 seconds, and 72°C for 15 seconds, with fluorescence detection at 522 nm. Relative gene expression was calculated using the 2-ΔΔCt method [95].

### Mass spectrometry-based Proteomics

Infected primary CD4^+^ T cells were harvested at 24 hpi and 48 hpi, washed with PBS, and centrifuged at 150 x g for 10 min to remove dead cells. 600000 cells were lysed in 50 µl proteomics lysis buffer (50 mM Tris pH 7.4, 250 mM NaCl, 25 mM EDTA, 1% NP-40, protease inhibitor) and stored at -20 °C until LC-MS/MS-based analysis (see below). In parallel, infection rate was assessed by flow cytometry. Proteins were proteolysed with LysC and trypsin with filter-aided sample preparation procedure (FASP) as described [96, 97]. Acidified eluted peptides were analyzed on a QExactive HF mass spectrometer (Thermo Fisher Scientific) online coupled to a UItimate 3000 RSLC nano-HPLC (Dionex). Samples were automatically injected and loaded onto the C18 trap cartridge and after 5 min eluted and separated on the C18 analytical column (Acquity UPLC M-Class HSS T3 Column, 1.8 μm, 75 μm x 250 mm; Waters) by a 90 min non-linear acetonitrile gradient at a flow rate of 250 nl/min. Data were recorded in data-independent mode (DIA) combining profile precursor spectra from 300 to 1,650 mass-to-charge ratio at 120,000 resolution with an automatic gain control (AGC) target of 3e6 and a maximum injection time of 120 ms, followed by fragmentation spectra covering 37 variable windows spanning from 300 to 1,650 m/z, each at 30,000 resolution with an AGC target of 3e6 and a normalized collision energy of 28.

The DIA LC−MS/MS data sets were analyzed using Spectronaut (Version 10, Biognosys, Schlieren, Switzerland) using an in-house spectral library based on data-dependent acquisition data from the same instrument searched against the SwissProt (Human) Release 2017_02, 20237 sequences with default settings. Only proteotypic peptides were included for protein quantification applying summed precursor quantities based on MS2 area quantity. Identifications were filtered with 1% false discovery rate on peptide and protein level. The normalised abundances were exported and processed as detailed in Supplemental Table S1 to calculate log2-fold modulations and according p-values using standard T-tests comparing each condition.

### Software and analyses

Flow cytometric analyses were performed using FlowLogic (v8.3). Absorbance data (for p24 quantification) was processed with ICE (version 1.0.9.8.) and imaging analysis was performed using Incucyte Base Analysis Software (v2022B). RT-qPCR data were processed using the LightCycler 480 Software (v1.5.1.62). Statistical analyses were performed using GraphPad Prism (v10.1.1) and Microsoft Excel. Figures were generated using CorelDRAW 2024 (v25.0.0.230) and GraphPad Prism (v10.1.1).

## Consent for publication

All authors gave their consent to publish. All authors read and approved the final manuscript.

## Availability of data and material

All data generated and analyzed during this study are included in this published manuscript.

## Authors’ contributions

J.L., A.D. and M.S. designed experiments. J.L. performed most of the experiments supported by A.D., C.A.V.-T., B.M., R.B. and N.W.. J.L., A.D., P.B., S.H. and M.S analyzed the data. S.H. and M.S. contributed reagents, funding and analysis tools. MS conceived the overall study. J.L. and M.S. prepared the first manuscript draft. All authors read and approved the final manuscript.

## Acknowledgements

We thank Éric A. Cohen (Montreal, Canada) for providing pWPI-based Vpr lentiviral expression plasmids, Frank Kirchhoff (Ulm, Germany) for providing the expression plasmids harboring primary vpr alleles and Daniel Sauter (Tübingen, Germany) for providing the TCF7-HA plasmid and the anti-β-catenin antibody. We are grateful to the team of the Transfusion Medicine Tübingen (ZKT Tübingen gGmbH, Tübingen, Germany) for continuous support in providing buffy coats. We thank Philip Bucher and Josef Leibold (Tübingen, Germany) for assistance with deciding on the subset markers.

---

**Supplementary Figure S1.**
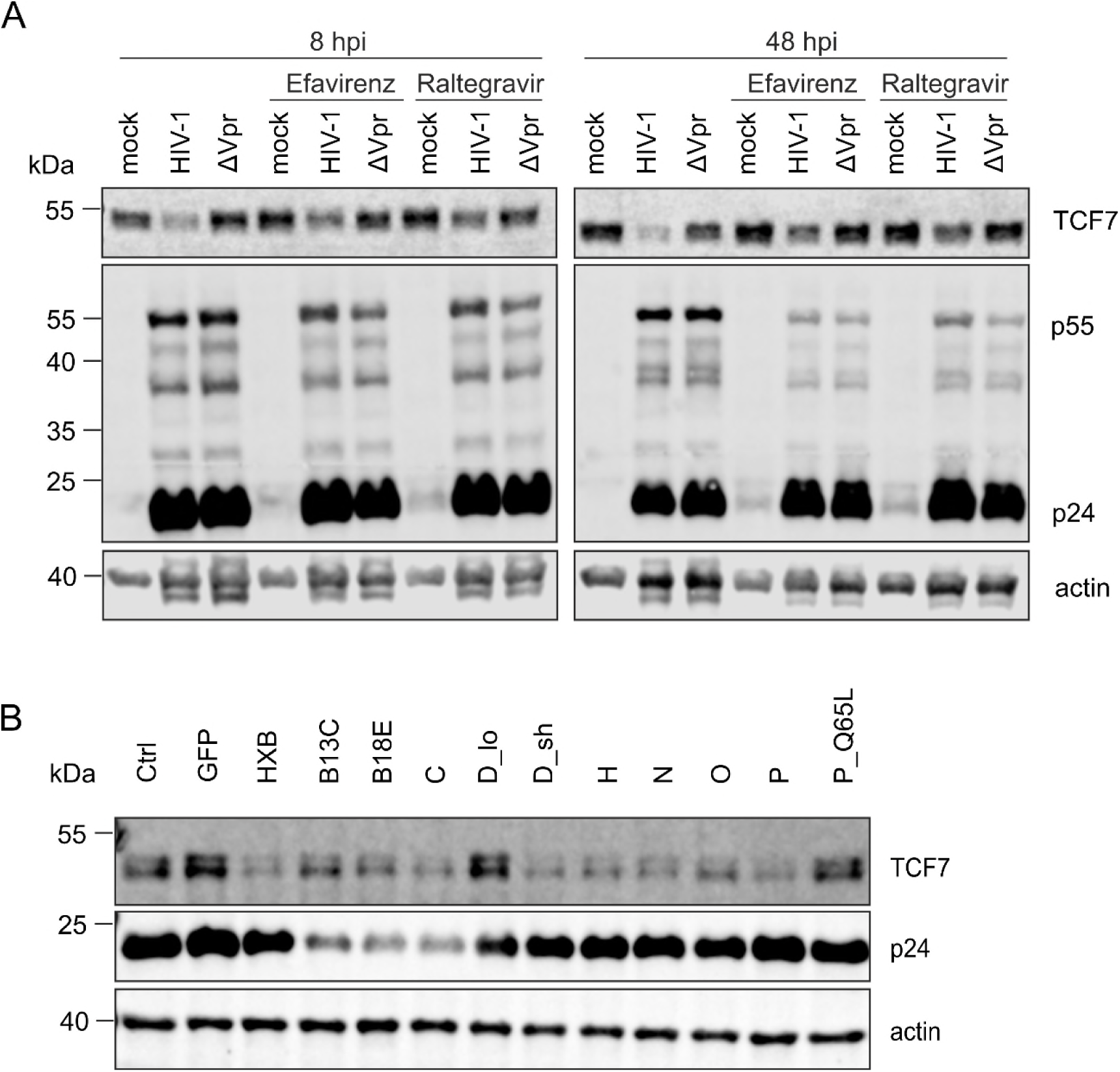
Incoming Vpr is sufficient to modulate TCF7 in CD4^+^ T cells. (A) Primary human CD4^+^ T cells were infected with HIV-1 or ΔVpr. Efavirenz (100 nM), Raltegravir (250 nM), or DMSO as a vehicle control was added at 2 hpi. Cells were lysed at 8 and 48 hpi and subjected to Western blot analysis of TCF7, HIV-1 p55, and HIV-1 p24. Actin was used as a loading control. Representative results from two biological replicates are shown. (B) SupT1 CD4^+^ T cells were transduced with lentiviral particles expressing Vpr alleles derived from different HIV- 1 subtypes. A Q65L mutant of the HIV-1 group P Vpr allele and an extended version of the HIV-1 group M subtype D Vpr allele (D_lo) were additionally included. “Ctrl” indicates cells transduced with virus-like particles containing no lentiviral genome, whereas “GFP” indicates cells transduced with virus-like particles containing a lentiviral genome expressing GFP only. Cells were lysed at 48 hpi and subjected to Western blot analysis of incoming Vpr and TCF7. Actin was used as a loading control. Representative blot from four biological replicates is shown.

**Supplementary Figure S2.**
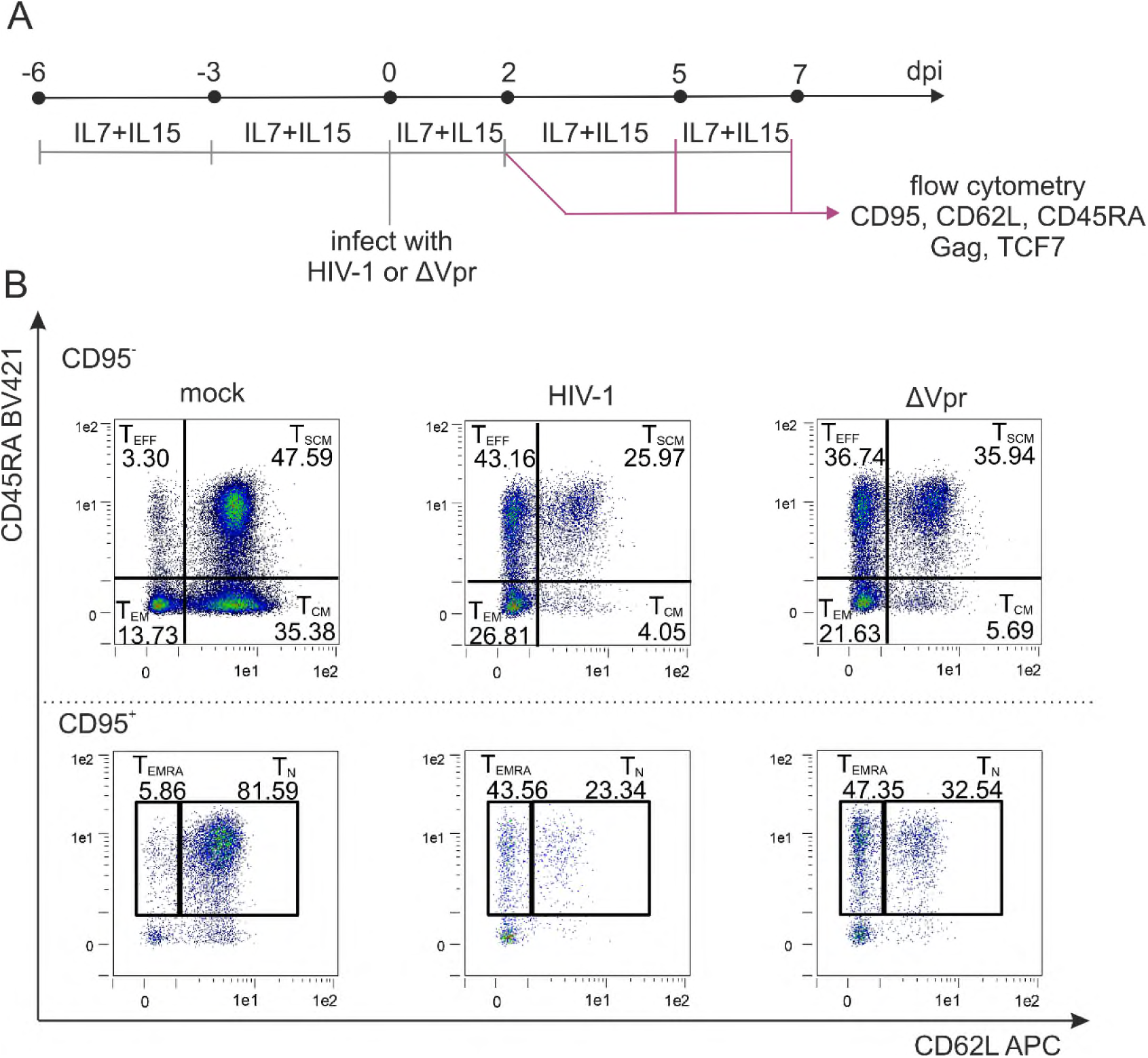
Experimental setup and gating strategy for the longitudinal analysis of primary CD4^+^ T-cell subsets. (A) Schematic representation of the experimental setup. Primary CD4^+^ T cells were isolated and maintained with IL-7 and IL-15 (20 ng/mL each) for 6 days before infection. Cells were infected with HIV-1 or HIV-1 ΔVpr and harvested at 2, 5, and 7 dpi for flow-cytometric assessment of CD95, CD62L, CD45RA, TCF7, and HIV-1 p24 expression. Cells received fresh medium supplemented with IL-7 and IL-15 every 2–3 days. (B) Representative flow-cytometry plots from eight biological replicates showing the different T-cell subsets at 5 dpi. CD62L and CD45RA expression are shown within previously gated CD95^−^ cells, defining T_EFF_, T_SCM_, T_CM_, and T_EM_ cells, and within CD95^+^ cells, defining T_EMRA_ and T_N_ cells.

**Supplementary Figure S3.**
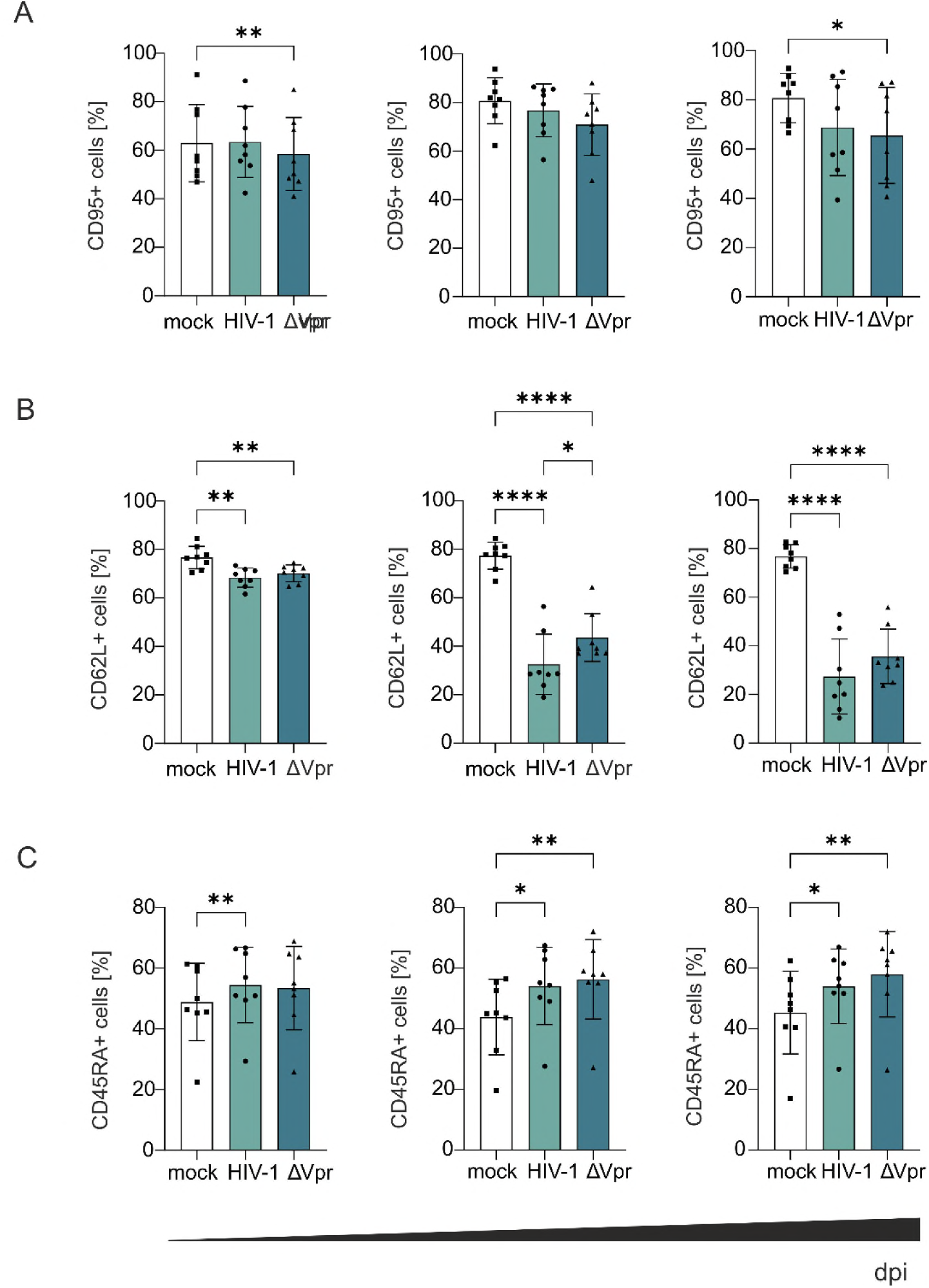
Modulation of CD95, CD62L, and CD45RA expression following HIV-1 infection. Primary CD4^+^ T cells maintained with IL-7 and IL-15 (20 ng/mL each) were infected with HIV-1 or ΔVpr. Surface expression of (A) CD95, (B) CD62L, and (C) CD45RA was determined between 2 and 7 dpi. Data are shown as the mean ± SD of eight biological replicates. Statistical significance was determined using paired one-way ANOVA with Tukey’s multiple-comparison test for CD95 and CD62L and the Friedman test for CD45RA. *p < 0.05; **p < 0.01; ****p < 0.0001.

**Supplementary Figure S4.**
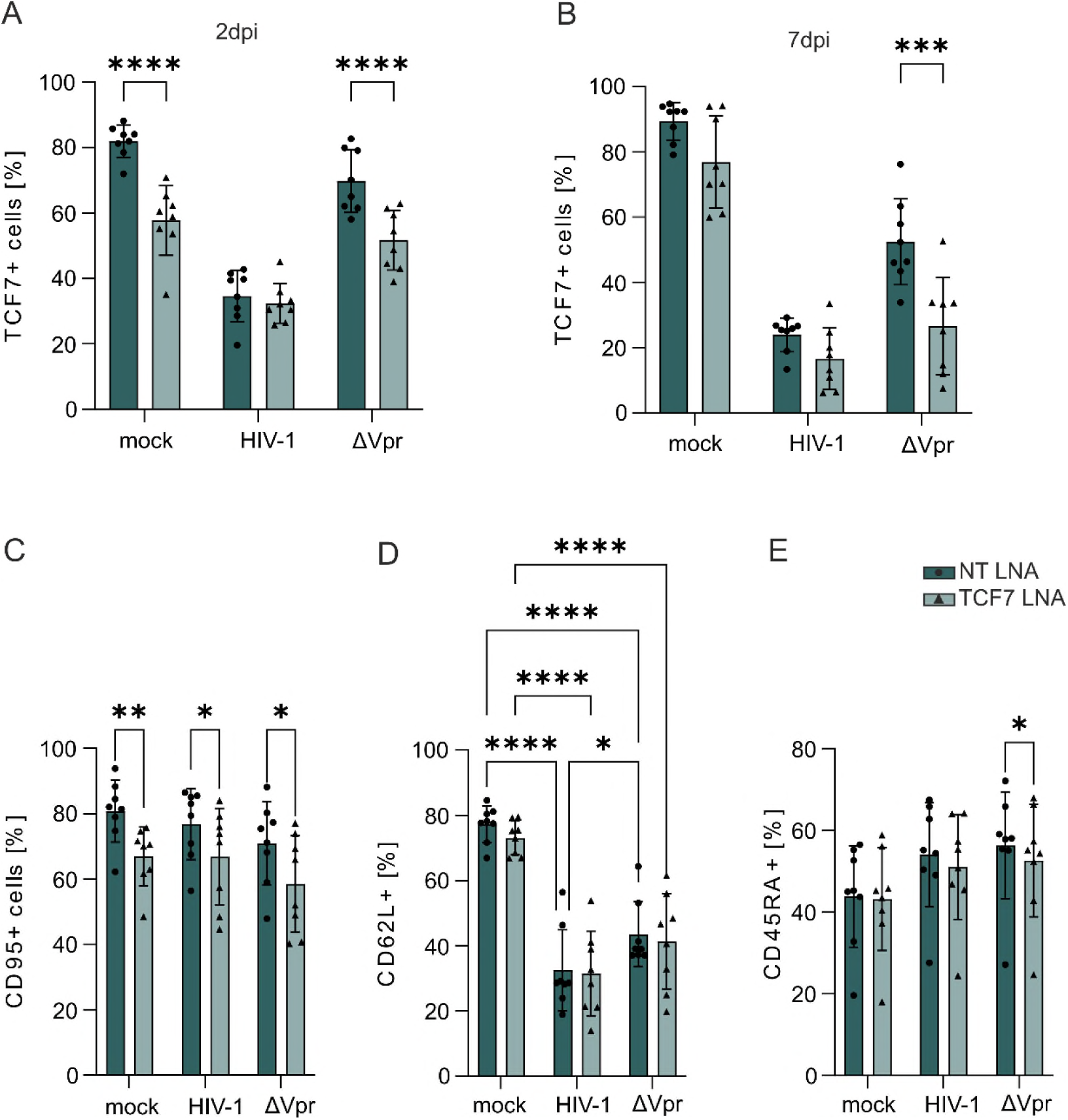
Effects of TCF7 knockdown on TCF7, CD95, CD62L, and CD45RA expression. Primary CD4^+^ T cells were stimulated with IL-7 and IL-15 (20 ng/mL each) and treated for 6 days with 6 µM non-targeting (NT) LNAs or with 3 µM each of two TCF7-targeting LNAs. On day 3, the medium was replaced with fresh medium supplemented with 20ng/ml IL-7 and IL-15, and the LNAs were readministered. Cells were infected with HIV-1 or ΔVpr and harvested at 2, 5, and 7 dpi for flow-cytometry based assessment of CD95, CD62L, CD45RA, TCF7, and HIV-1 p24 expression. TCF7 knockdown efficiency was determined by intracellular flow-cytometric staining of TCF7 at (A) 2 dpi and (B) 7 dpi. Surface expression of (C) CD95, (D) CD62L and (E) CD45RA was determined at 5 dpi. n=8 (Mean ± SD). Statistical significance was determined using a paired two-way ANOVA with Šídák’s multiple-comparison test. *p < 0.05; **p < 0.01; ****p < 0.0001.

**Supplementary Figure S5.**
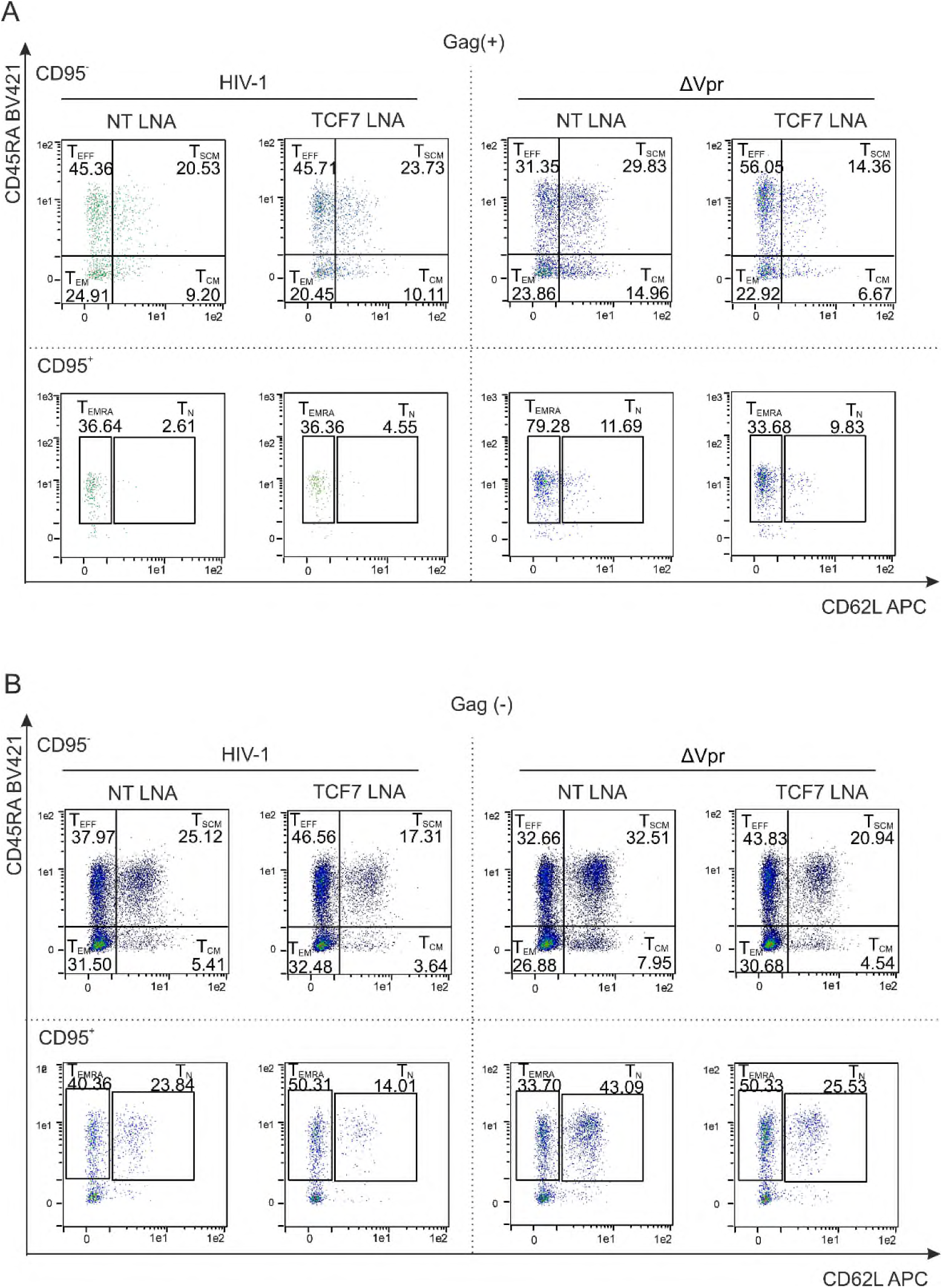
Gating strategy for T-cell subsets within infected and bystander cells. Flow-cytometry plots showing the different T-cell subsets at 5 dpi. CD62L and CD45RA expression are shown within previously gated CD95^−^ cells, defining T_EFF_, T_SCM_, T_CM_, and T_EM_ cells, and within CD95^+^ cells, defining T_EMRA_ and T_N_ cells. Representative plots from n=8 are shown for (A) infected Gag (+) cells and (B) bystander Gag (−) cells.

**Supplementary Figure S6.**
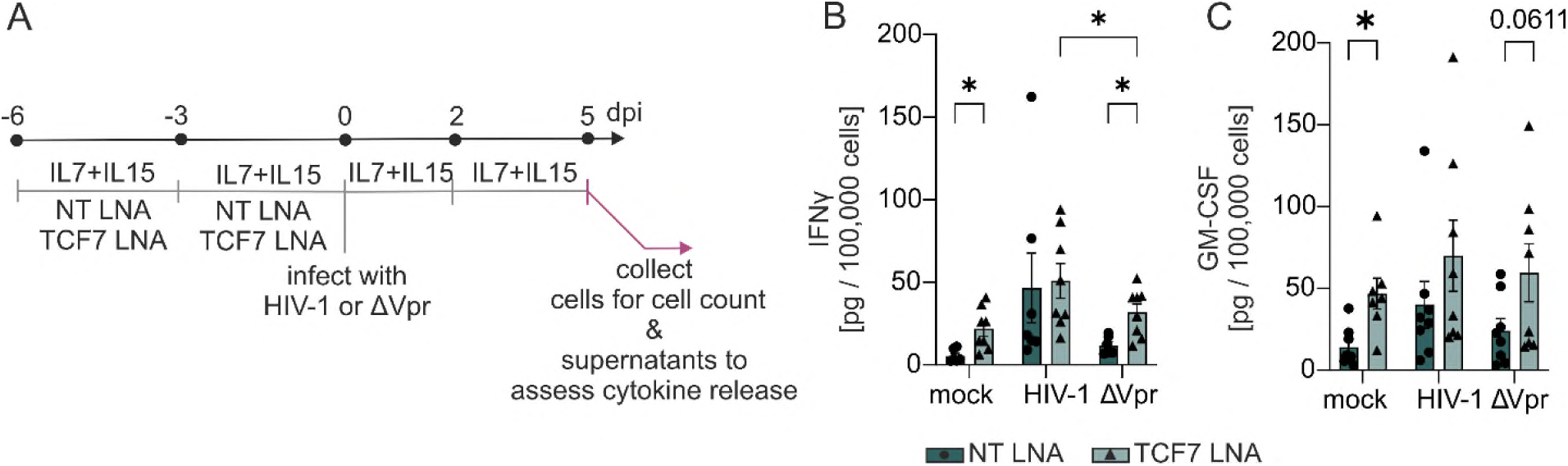
Cytokine release following HIV-1 infection of primary CD4^+^ T cells. (A) Schematic representation of the experimental setup. Primary CD4^+^ T cells were isolated and maintained in the presence of IL-7 and IL-15 (20 ng/mL each) for 6 days before infection. Cells were infected with HIV-1 or ΔVpr, and cells and culture supernatants were harvested at 5 dpi for assessment of cell numbers and cytokine release. Cytokine concentrations in the culture supernatants were determined using LEGENDplex. (B) Release of IFN-γ and (C) GM-CSF into the culture supernatant are shown. n = 6–8 (Mean ± SEM) for IFN-γ, n = 7-8 (Mean ± SEM) for GM-CSF. Values below the lower limit of detection were excluded. Statistical significance was determined using a paired mixed-effects analysis with Šídák’s multiple-comparison test. *p < 0.05.

